# SARS-CoV-2 EndoU-ribonuclease regulates RNA recombination and impacts viral fitness

**DOI:** 10.1101/2024.11.11.622995

**Authors:** Yiyang Zhou, Yani P. Ahearn, Kumari G. Lokugamage, R. Elias Alvarado, Leah K. Estes, William M. Meyers, Alyssa M. McLeland, Angelica L. Morgan, Jordan T. Murray, David H. Walker, Bryan A. Johnson, Andrew L. Routh, Vineet D. Menachery

**Author notes:** **Corresponding Author:** Andrew L. Routh, Vineet D. Menachery, **Email:**. Co-First Authors (Yiyang Zhou, Yani P. Ahearn). Co-Senior Authors (Andrew L. Routh, Vineet D. Menachery).

## Abstract

Coronaviruses (CoVs) maintain large RNA genomes that frequently undergoes mutations and recombination, contributing to their evolution and emergence. In this study, we find that SARS-CoV-2 has greater RNA recombination frequency than other human CoVs. In addition, coronavirus RNA recombination primarily occurs at uridine (U)-enriched RNA sequences. Therefore, we next evaluated the role of SARS-CoV-2 NSP15, a viral endonuclease that targets uridines (EndoU), in RNA recombination and virus infection. Using a catalytically inactivated EndoU mutant (NSP15^H234A^), we observed attenuated viral replication *in vitro* and *in vivo*. However, the loss of EndoU activity also dysregulated inflammation resulting in similar disease *in vivo* despite reduced viral loads. Next-generation sequencing (NGS) demonstrated that loss of EndoU activity disrupts SARS-CoV-2 RNA recombination by reducing viral sub-genomic message but increasing recombination events that contribute to defective viral genomes (DVGs). Overall, the study demonstrates that NSP15 plays a critical role in regulating RNA recombination and SARS-CoV-2 pathogenesis.

## Introduction

The emergence of the SARS-CoV-2 in 2019 resulted in a global pandemic with unprecedented economic disruption and 700 million cases worldwide [1, 2]. While initial efforts to quell the outbreak focused on vaccination [3], the development of SARS-CoV-2 variants of concern (VoCs) demonstrated the ability of the virus to evolve and evade host immunity [4, 5]. As a result, “herd” immunity to COVID-19 has rendered a less deadly, but a still quite infectious and transmissible SARS-CoV-2. Importantly, the continued evolution of SARS-CoV-2 suggests that most people will face multiple infections and potential long-term complications including numerous manifestations of long COVID [6, 7].

Mutation and recombination are the main drivers of CoV evolution. While employing an error prone polymerase like other RNA viruses, CoVs have a significantly lower mutation rate governed by a proofreading viral 3’ exonuclease [8, 9]. Yet, the sheer number of SARS-CoV-2 infections worldwide has led to accumulation of advantageous mutations and evolution of variants [5]. RNA recombination offers a second mechanism for adaptation, shuffling of chunks of genetic sequence within and between virus strains [10]. Importantly, recombination is required for the CoV lifecycle including its generation of sub-genomic messenger RNA (sgmRNA) from discontinuous genome segments [11]. In addition, genetic and experimental analyses reveal extensive recombination between virus strains of the CoV families [12, 13]. Giving rise to hybrid and novel strains, these recombinant viruses may be the key to viral emergence and immune evasion. Finally, RNA recombination gives rise to defective viral genomes (DVGs) which play a complex and still unresolved role in engaging host immunity following infection [14]. These activities highlight the importance of RNA recombination to CoV infection and identify the need to better understand its underpinnings.

Despite playing a critical role, CoV RNA recombination is still poorly understood. Prior to the development of next generation sequencing (NGS), analysis of recombination was severely limited and difficult to study [15]. Even early NGS approaches have complicated analysis with PCR duplication, error rate, and other sequencing artifacts reducing confidence in potential findings. However, as the SARS-CoV-2 pandemic continued, novel techniques and approaches have allowed further insights into CoV recombination. Initial work by our group found that SARS-CoV-2 is more recombinogenic than MERS-CoV [15]. The work has also implicated viral exonuclease NSP14 in playing a role in promoting recombination in mouse hepatitis virus. Similarly, recombination has been reported to induce significant DVG production following SARS-CoV-2 infection driving host immune responses [14]. Notably, SARS-CoV-2 was also shown to primarily recombine at uridine rich sequences [15]; these U-rich tracts are potential targets for CoV NSP15, a highly conserved viral endonuclease targeting uridines (EndoU) [16–19]. Prior work has shown CoV EndoU plays a critical role in preventing host sensor recognition by cleaving viral RNA and preventing interferon responses [17, 18, 20–22]. Given that RNA recombination junctions primarily occur at uridine-rich tracts, NSP15 may contribute to CoV RNA recombination.

In this study, we explore RNA recombination in the context of SARS-CoV-2 and other human CoVs. Using a refined analysis pipeline, we demonstrate that SARS-CoV-2 RNA is more recombinogenic than other human CoVs. We also show that RNA recombination occurs primarily at uridine-enriched tracts across each of the HCoVs. Mechanistically, the uridine-rich sequence at the RNA recombination junctions suggested a role for CoV endoribonuclease NSP15. Therefore, we generated a catalytically inactive NSP15 mutant (NSP15^H234A^). NSP15^H234A^ shows attenuated viral replication *in vitro* and *in vivo*, but similar *in vivo* pathogenesis of wild-type (WT) infection, which is driven by augmented host responses characterized by both antiviral activity and inflammation mediated tissue damage. Surprisingly, loss of NSP15 activity increased recombination events *in vitro* including deletions and micro-deletions; yet, NSP15^H234A^ also had reduced viral subgenomic mRNA formation. *In vivo,* NSP15^H234A^ continued to show reduced viral subgenomic mRNA formation. In addition, NSP15^H234A^ infected animals contain a viral population with reduced diversity but strong selection of particular defective viral genome populations. Overall, our results highlight a critical role for NSP15 in modulating different RNA recombination (facilitating sgmRNA formation but antagonizing DVGs), which contribute to the development of viral infection, pathogenesis, and host immune responses.

## Results

### Increased RNA recombination in SARS-CoV-2 compared to other human coronaviruses

Our prior studies suggested that SARS-CoV-2 RNA was more recombinogenic than MERS-CoV RNA [15]. To determine if SARS-CoV-2 RNA produced greater recombination frequency than other human coronaviruses (HCoVs), we conducted additional experiments with SARS-CoV-2 and two common cold HCoVs strains, HCoV-OC43 and HCoV-229E. Briefly, appropriate cell lines (Vero E6, HUH7, and HCT8) were infected with SARS-CoV-2, HCoV-229E, or HCoV-OC43 at low MOI. When significant cytopathic effect (>40%) was observed, total cellular RNA was collected and next-generation sequencing (NGS) libraries were constructed using the random-primed ClickSeq approach [23]. NGS reads were processed and aligned to corresponding virus genomes, and analysis was conducted with bioinformatic pipeline “*ViReMa (Virus Recombination Mapper)*” to map the distribution of RNA recombination events (**Fig. 1a**) [24, 25]. In addition, our previously published MERS-CoV data [15] were reanalyzed using the same bioinformatic pipeline to facilitate comparisons.

**Figure 1.**
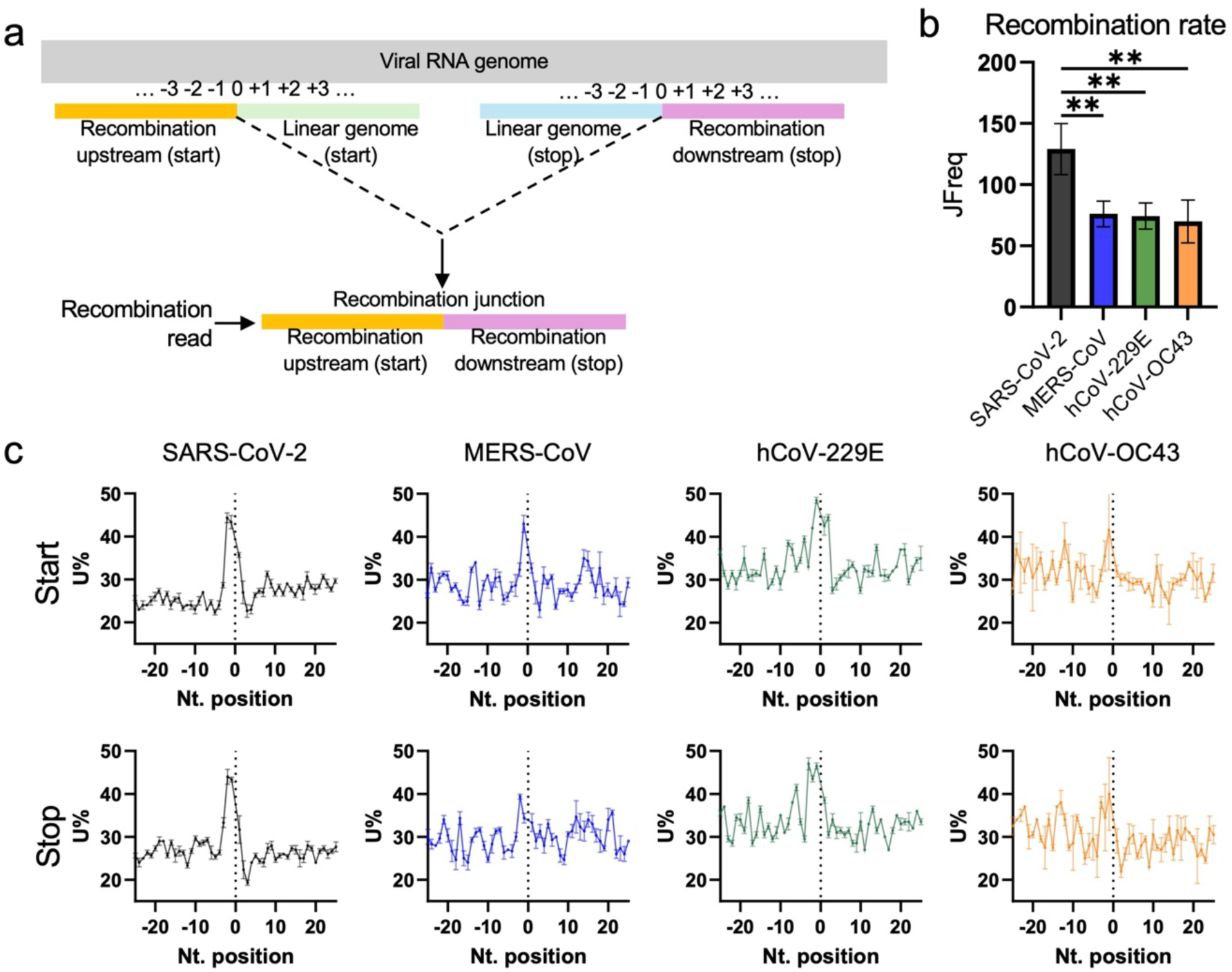
Human coronavirus RNA recombination favors U-rich tracks. (a) Schematic of RNA recombination reads that consist of gaps in linear genome, and the nucleotide positions flanking the recombination start (donor) and stop (acceptor) sites. (b) Cell culture infected with human coronaviruses and sequenced with random primers. SARS-CoV-2 RNA showed higher recombination tendency than other human coronaviruses. “JFreq” (junction frequency) measures the number of recombination events per 104 sequenced virus reads. (c)The RNA recombination of human coronaviruses utilizes U-rich sequences flanking the recombination junctions. Error bar: standard deviation. **: P value < 0.01 (one-way ANOVA, α = 0.05, N=3 for SARS-CoV-2 and MERS-CoV; N=2 for hCoV-229E and hCoV-OC43).

Analyzing junction frequency (JFreq, the number of ViReMa-detected recombination junctions per 10^4^ mapped viral reads [15]), revealed a significantly higher frequency of recombination for SARS-CoV-2 than MERS-CoV, HCoV-229E, and HCoV-OC43 (**Fig. 1b**). While the other HCoVs had comparable recombination events that hovered at JFreq of 70-76, SARS-CoV-2 had a ∼1.8 fold greater JFreq of ∼125. These results demonstrate that SARS-CoV-2 RNA is more recombinogenic than other human CoVs, consistent with our earlier studies with MERS-CoV [15].

### Human Coronavirus RNA recombination occurs most frequently at U-rich tracts

Our previous studies found that SARS-CoV-2 RNA recombination was enriched at uridine-rich tracks flanking the ‘start’ and ‘stop’ sites of recombination junctions, both in cell culture and from human clinical specimens [15, 26]. Here, we evaluated whether the U-favored RNA recombination applies to other human coronaviruses. The uridine and other nucleotide percentages were calculated at each upstream (-25 to -1) or downstream (+1 to +25) positions of the recombination junction, as well as the linear genome positions (**Fig. 1a**). Each recombination event was also weighted by abundance to provide a representative understanding of the nucleotide frequency at each position. However, contrasting the prior approach [15], sub-genomic mRNA (sgmRNA) events were excluded due to their predominance and putatively different recombination mechanisms. We observed distinct peaks of uridine percentage (U%) flanking recombination junctions in all 4 HCoV infections (**Fig. 1c**), while no robust trends were observed in the other nucleotides (**Extended Data Fig. 1**). Our results indicate that the RNA recombination of all four HCoVs are most frequent at uridine(U)-rich sequences near both start (donor) sites and stop (acceptor) sites of RNA recombination and that the U-enriched RNA recombination is not unique to SARS-CoV-2 but applies to the other HCoVs tested.

### Loss of EndoU activity attenuates SARS-CoV-2 replication *in vitro*

Given the propensity for CoV recombination to occur in uridine-enriched RNA tracts, we next focused on CoV non-structural protein 15 (NSP15), an endoribonuclease that cleaves RNA at uridine rich sites (EndoU) [17]. NSP15 is known to play a key role in evading type I interferon (IFN) by targeting viral RNA for cleavage and disrupting recognition by host sensors [17, 20]. Importantly, NSP15 activity is maintained across the entire CoV family, and the active site residues are conserved (**Fig. 2a & b**) [27]. While established to play a role in CoV infection and immune evasion, we sought to determine if NSP15 EndoU activity impacts SARS-CoV-2 viral RNA recombination.

**Figure 2.**
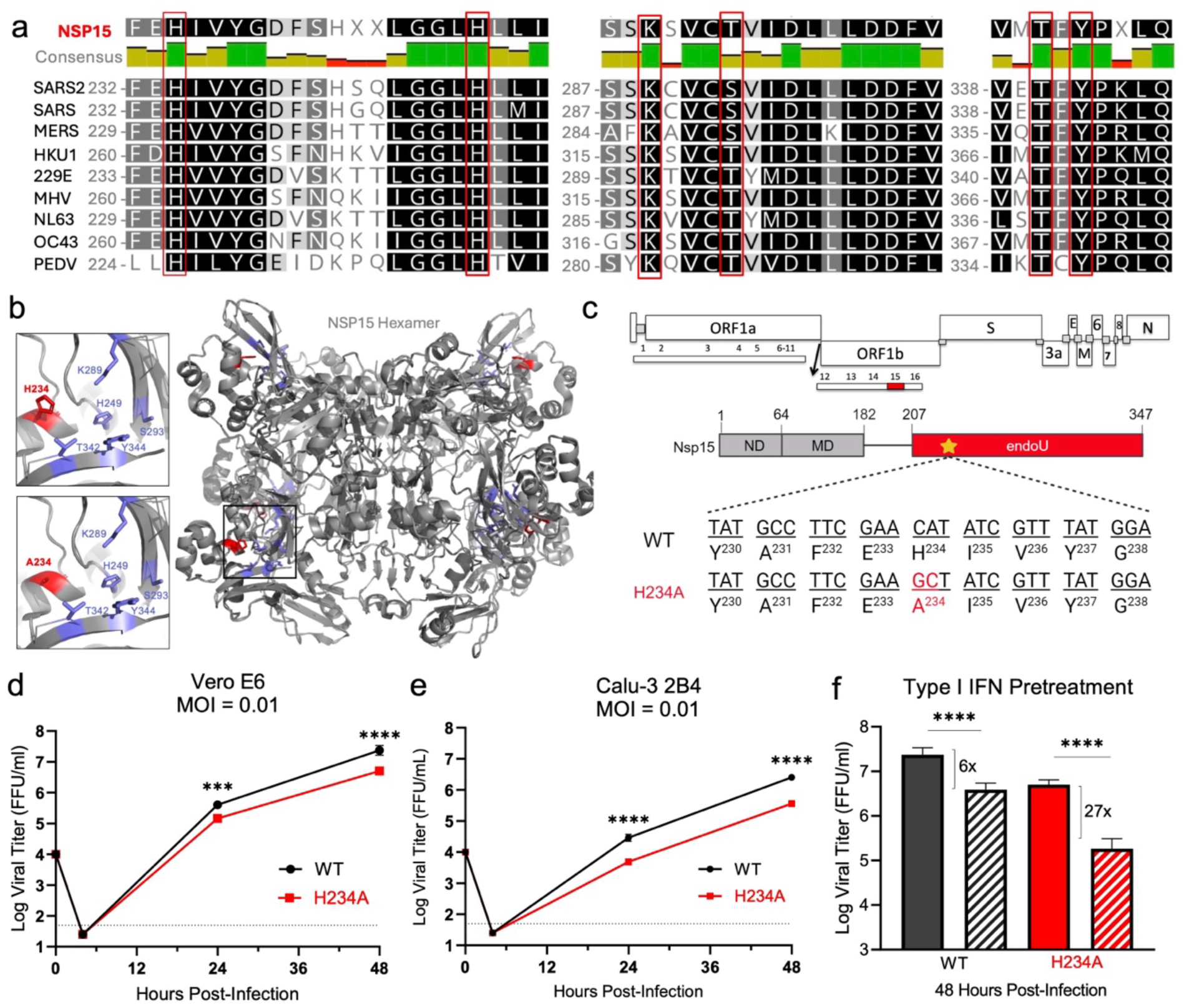
SARS-CoV-2 Nsp15 mutant (H234A) in vitro characterization. (a) Sequence alignment of Nsp15 endoribonuclease domain from different coronaviruses. (b) SARS-CoV-2 Nsp15 hexamer (grey) with catalytic amino acid residues labeled (blue). The histidine-to-alanine mutation at amino acid position 234 is in red. (c) Schematic of the Nsp15 structure showing the N-terminal domain (ND), middle domain (MD), and endoribonuclease domain (endoU). Nucleotides in red represent the 2-bp substitution in the H234A mutant. (d) Viral replication in Vero E6 cells infected with WT (black) or H234A (red) at MOI = 0.01 (n=6 from two experiments with three biological replicates each). (e) Viral replication of Calu-3 2B4 cells infected with WT (black) or H234A (red) at MOI = 0.01 (n=6 from two experiments with three biological replicates each). (f) Vero E6 cells were treated with control (solid) or 100 U type I interferon (IFN) (hashed) 16 hours prior to infection with WT (black) or H234A (red) at MOI = 0.01. Viral replication was measured at 48 hours post infection (n=6 from two experiments with three biological replicates each). The fold change relative to control is shown in brackets for each virus. Data are presented as mean ± SD. Statistical analysis was measured using a two-tailed Student’s t-test. **** P < 0.001.

While NSP15 deletion mutants are not viable, catalytically inactive mutants have been recovered and characterized in several CoVs including mouse-hepatitis virus (MHV) and MERS-CoV [20, 28]. In this study, we took a similar approach by targeting amino acid H234 to ablate catalytic activity as previously described [19] (**Fig. 2a-c**). Using our established SARS-CoV-2 reverse genetic system [29, 30], we generated a NSP15 mutant (NSP15^H234A^) capable of robust replication without significant changes in plaque morphology. Following inoculation of Vero E6 cells at MOI 0.01, the NSP15^H234A^ exhibited modest attenuation (∼ 0.5 log) in viral replication at both 24- and 48-hours post-infection (HPI) relative to the WT SARS-CoV-2 virus (**Fig. 2d**). These results suggest that the NSP15^H234A^ had a small impact on the viral replication capacity. We then examined viral replication in Calu-3 2B4 cells, an interferon (IFN)-responsive human respiratory cell line. We observed a more robust reduction (∼ 1 log) in viral replication of the NSP15^H234A^ at both 24 and 48 HPI (**Fig. 2e**). Taken together, our data demonstrate that the disruption of the catalytic endoU domain in NSP15 attenuates viral replication *in vitro*.

### NSP15^H234A^ has increased sensitivity to type I interferon

Prior studies demonstrate the importance of CoV NSP15 in controlling the type I IFN response following infection [17, 20]. To examine IFN sensitivity of NSP15^H234A^, we pretreated Vero E6 cells with 100 units of universal type I IFN and infected at MOI 0.01. Compared to untreated cells, WT SARS-CoV-2 had a modest reduction in viral replication (∼6 fold) following type I IFN pretreatment, consistent with previous findings (**Fig. 2f**) [31]. In contrast, the NSP15 mutant virus had a 27-fold reduction in viral titers (**Fig. 2f**). These results indicate that the NSP15^H234A^ is more sensitive to type I IFN responses than WT SARS-CoV-2. These results are consistent with findings from MHV and MERS-CoV [20, 28].

### NSP15^H234A^ attenuates viral replication, but not disease *in vivo*

Having established attenuation *in vitro*, we next evaluated the NSP15^H234A^ *in vivo* using the Golden Syrian Hamster model of infection [32, 33]. Briefly, three-to-four-week-old golden Syrian hamsters were challenged with either WT SARS-CoV-2 or NSP15^H234A^ at 10^5^ focus forming units (FFU) and monitored for weight loss and disease over a 7-day time course (**Fig. 3a**). At 2, 4, and 7 days post-infection, cohorts of animals were nasal washed under anesthesia, subsequently euthanized, and lung tissues collected for further analyses of viral titers and histopathology. Surprisingly, hamsters infected with NSP15^H234A^exhibited similar weight loss and disease as WT-infected animals (**Fig. 3b**). These results contrast *in vitro* findings and indicate that NSP15^H234A^ maintains the capacity to cause significant disease *in vivo*.

**Figure 3.**
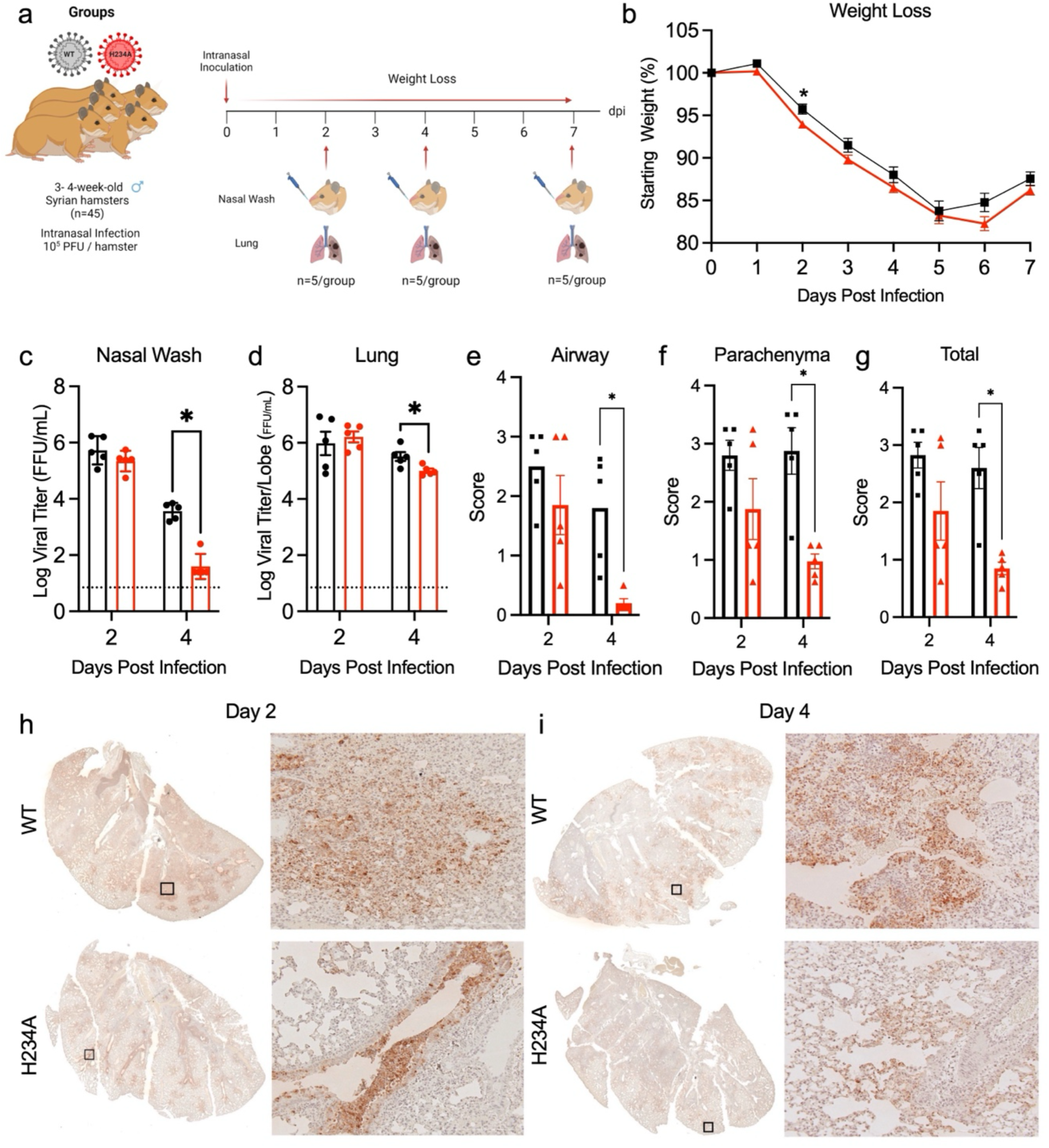
In vivo characterization of NSP15^H234A^. (a) Schematic of golden Syrian hamster infection with WT or H234A SARS-CoV-2. Three-to-four-week-old golden Syrian hamsters were intranasally infected with 10^5^ plaque forming units (PFU) of WT or H234A and monitored for weight changes and signs of disease over a 7 day time course. Hamsters were nasal washed, and lung lobes were collected at days 2, 4, and 7 post infection for viral titers and histopathology analyses. (b) Percent body weight change from starting weight for WT or H234A infected groups. (c-d) Viral titers were measured for nasal wash (c) and lung (d) for animals infected with WT or H234A at day 2 and 4 post infection. ((e-g) Scores of viral antigen staining in e) airway, f) parenchyma, and g) total from hamster left lung infected with WT or H234A. (h-i) Representative images of viral antigen staining (nucleocapsid) of hamster lung sections at h) day 2 or i) day 4. Data are presented as mean ± SD. Statistical analysis was measured using a two-tailed Student’s t-test. **** P < 0.05.

Examining viral load in the nasal wash and lung, we observed that both WT and NSP15^H234A^ infected animals had similar viral titers at day 2 post infection (**Fig. 3c & d**). However, by day 4 post infection, we observed significant reductions in viral titers in the nasal wash (∼ 2 log) and lung (∼0.5 log) viral titers of the NSP15^H234A^ infected hamsters relative to WT-infected animals (**Fig. 3c & d**). Similarly, viral antigen staining in the lung also showed reduced infection in the NSP15^H234A^ infected animals as compared to WT controls (**Fig. 3e-i**). Airway, parenchyma, and overall lung antigen scoring showed a significant deficit in the NSP15^H234A^ compared to WT at day 4 with similar trends at day 2(**Fig. 3e-g**). While antigen distribution was similar, overall staining intensity and area were diminished in the mutant relative to WT (**Fig. 3 h & i).** Together, the viral titer data and antigen staining demonstrate attenuation of viral replication in NSP15^H234A^ despite significant weight loss following infection.

### Significant disease and damage observed following NSP15^H234A^ infection

Having established reduced viral loads and antigen staining in the lung, we further evaluated disease and damage in the lung of NSP15^H234A^ infected hamsters. Utilizing H&E staining, a certified pathologist examined lung sections from days 2 and 4 following WT and NSP15^H234A^ infection (**Extended Data Fig. 2a**). Despite differences in viral antigen staining, both WT and NSP15^H234A^ infected hamsters had significant immune infiltration and damage relative to mock (**Extended Data Fig. 2b-f**). At day 2, little pathology was observed with any groups, consistent with previous studies of SARS-CoV-2 in hamsters [33–35] (**Extended Data Fig. 2c & d**). However, both WT and NSP15^H234A^ viruses had severe disease at day 4 characterized by bronchiolitis, interstitial pneumonia, vasculitis, and alveolar cytopathology (**Extended Data Fig. 2e & f**). Importantly, the disease and damage in the H&E scores reflected a massive immune infiltrate and damage in the NSP15 mutant infection despite reduced viral loads. Together, the inflammation and immune infiltration likely lead to lung damage resulting in the similar weight loss and disease observed between mutant and WT virus infected animals.

### NSP15^H234A^ mutant induced augmented host immune responses *in vivo*

Coronavirus NSP15 has been shown to play an important role in cleaving viral RNA and preventing recognition by host innate immune sensors [16, 17, 36–38]. To evaluate changes in host responses, we profiled the transcriptomes of WT and NSP15^H234A^ mutant at days 2 and 4 post infection. Total cellular RNA from hamster lung tissues were sequenced with Poly(A)-ClickSeq as previously described [39, 40]. By mapping reads to the *Mesocricetus auratus* (Golden Hamster) reference genome, we obtained gene counts across 15606 annotated and unknown genes. Our results show divergent transcriptomic profiles between WT and NSP15^H234A^, especially at day 2 (**Fig. 4a**, red box). By day 4, this divergence was mostly lost with the NSP15^H234A^-infected lungs having a similar gene expression profile to WT infected lungs. Principal component assay (PCA) confirms that the host responses against NSP15^H234A^ diverged from WT at day 2, while day 4 infections and PBS mock controls clustered respectively (**Fig. 4b**).

**Figure 4.**
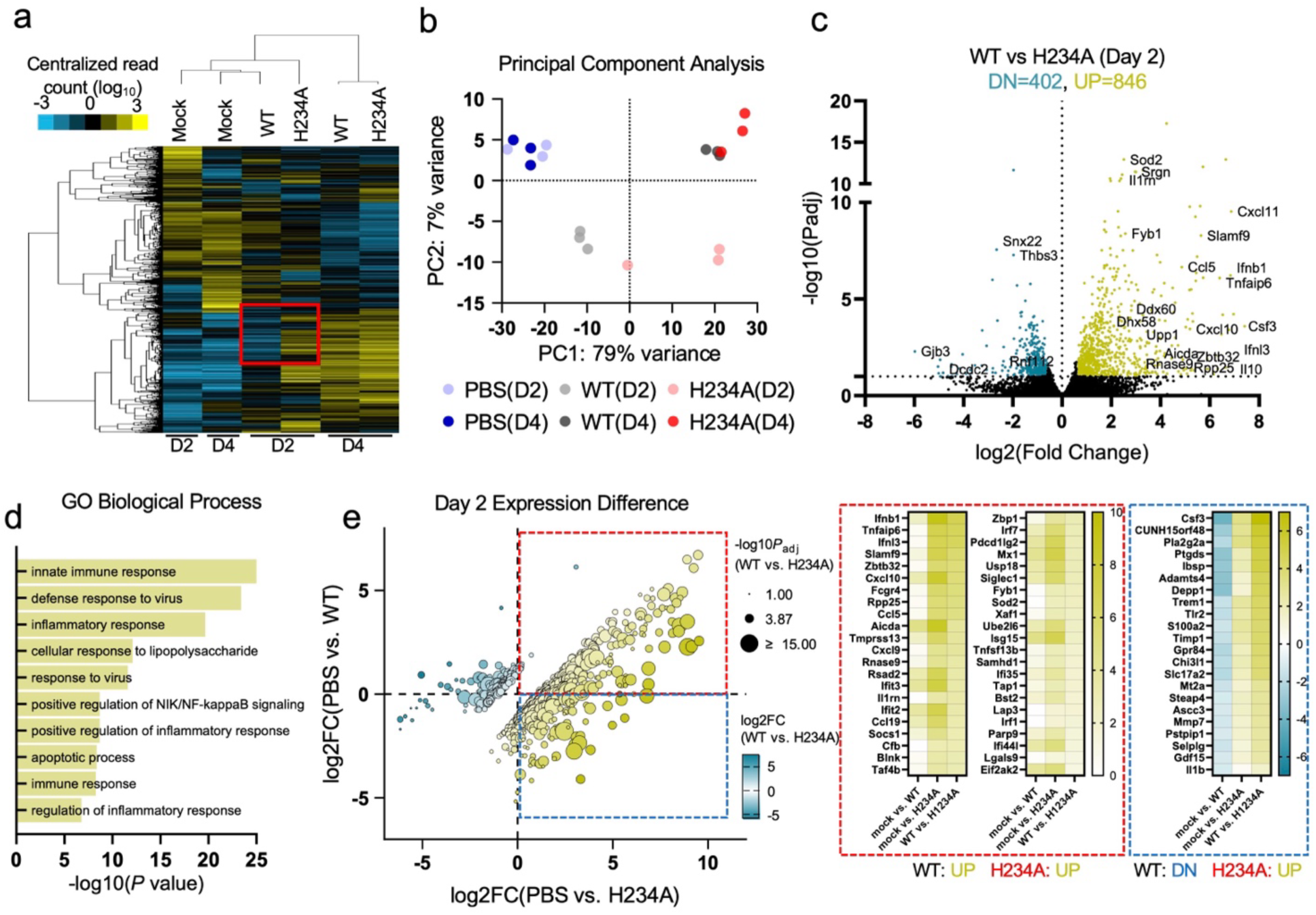
NSP15^H234A^ resulted in elevated host immune responses during early infection. (a) Hierarchical gene clusters based on average read count of each gene. H234A and WT showed distinctive gene expression profiles with particular clusters of genes (red box) at day 2 (D2) but not at day 4 (D4). (b) Principal component assay (PCA) showed that host responses against H234A progressed faster than WT at D2 but comparable at D4. (c) Up- and down-regulated genes between H234A and WT at D2 (padj0.585, N=3). (d) Top 10 gene ontology (GO) terms discovered from significantly (padj < 0.1, |log2FC| > 1, N=3) up- or downregulated genes (WT vs H234A) at D2. (e) At D2, WT and H234A showed both correlated and anti-corelated gene expression profiles in comparison to mock. Red box: genes that are up-regulated for both WT and H234A; blue box: genes that are down-regulated by WT but up-regulated by H234A; green box: genes that are up-regulated by WT but down-regulated by H234A.

Gene expression analyses (**Extended Data Fig. 3**) revealed 1266 differentially expressed genes between WT and H234A at day 2 (*p*_adj_ < 0.1, |fold change|>1.5); 864 with increased expression in H234A and 402 with decreased expression. In contrast, only 57 differentially expressed genes were observed at day 4 (*p*_adj_ < 0.1, |fold change|>1.5): 42 with increased expression in H234A and 15 with decreased expression (**Extended Data Fig. 3&4**). The results indicate that differential host responses occur between NSP15^H234A^ and WT infection at early times post infection.

We further scrutinized how NSP15^H234A^ induced different host response than WT at day 2 (**Fig. 4c**). Among the upregulated genes, we identified enrichment of immune modulatory genes (e.g., *Ifnb1, Tnfaip6, Ifnl3, Cxcl10*) as well as several genes known for their function in dsRNA-binding and RNA cleaving/modification (e.g., *Zbtb32, Rpp25, Aicda, Upp1, Ddx60,* and *Dhx58*). Our results suggest that the catalytically active NSP15 endonuclease activity suppresses the activation of these host immune factors. We subsequently conducted DAVID [41] gene ontology (GO) analysis using a curated list of 444 differentially upregulated genes and 171 differentially downregulated genes of H234A(D2) (*p*_adj_ < 0.1, |log2FC| > 1, relative to WT(D2)). The top 10 (ranked by *p*-value) GO terms in biological process, cellular component, and molecular function are presented (**Fig. 4d, Extended Data Fig. 5**). Examining the GO biological processes, several are associated with either innate immunity or response to viral infection (innate immune response, defense response to virus, cellular response to LPS, response to virus, immune response), suggesting loss of NSP15 activity is driving increased host sensing and type I interferon responses. However, several GO processes augmented in the NSP15^H234A^-infected animals are associated with inflammation, overactive immune responses, and apoptosis (inflammatory response, positive regulation of NIK/NF-kB Signaling, positive regulation of inflammatory response, apoptotic process). Together, the results indicate that NSP15^H234A^ induces a robust antiviral response controlling virus infection coupled with excessive inflammation driving immune mediated pathology.

To further understand the host responses, we compared the transcriptomics of WT and NSP15^H234A^ vs. PBS mock control (**Fig. 4e**). At day 2, most differentially expressed genes (*p*_adj_ < 0.1) between WT and NSP15^H234A^ showed a clear correlation, indicating that host responses to NSP15^H234A^ follow the same trajectory as WT, but elicit greater gene expression relative to WT. Examination of genes upregulated in both infections (**Fig. 4e**, red box) finds augmented antiviral factors in the NSP15^H234A^-infected animals including interferon genes (e.g., *Ifnb1, Ifnl3)* and interferon stimulated genes (e.g., *Rsad2, Ifit3, Ifit2, Mx1, Isg15, Bst2*). These results are consistent with the finding that the loss of NSP15 activity increased host sensing of viral RNA and augmented innate immune responses. At the same time, the cluster of genes upregulated in both infection groups also has several inflammatory markers (e.g., *Tnfaip6, Il1rn, Socs1, Tnfsf13b)* and cytokines (e.g., *Cxcl10, Ccl5, Cxcl9, Ccl19).* This demonstrates that the lack of NSP15 activity also elicits stronger inflammatory responses. In addition, the anti-correlatively expressed genes between WT and NSP15^H234A^ (suppressed in WT but induced in NSP15^H234A^, **Fig. 4e**, blue box) also have several factors associated with inflammation (e.g., *Csf3*, *Pla2g2a, Ptgds, IL1b, MMp7)*, suggesting that NSP15^H234A^ induces the expression of inflammatory genes which are typically suppressed in WT infection. Together, these results demonstrate that NSP15^H234A^ drives an augmented immune response, pairing antiviral activity with excessive inflammation and damage during early infection. This is also consistent with the *in vivo* observations stated above (**Fig. 3**), with the transcriptomic changes preceding pathology observations.

### NSP15^H234A^ increased RNA recombination but reduced sgmRNAs *in vitro*

Having established viral replication attenuation and altered host responses, we next evaluated changes in viral RNA recombination of WT and NSP15^H234A^. Total cellular RNA was collected from Vero E6 cells infected with WT or NSP15^H234A^ (**Fig. 5a**). NGS libraries were made using Tiled-ClickSeq (TCS) [26], an approach that uses >300 primers targeting the entire SARS-CoV-2 genome to provide sensitive detection and even coverage across the virus genome. TCS also allows for improved RNA recombination resolution [26] and better effectiveness than a random priming approach for sequencing SARS-CoV-2 genome from cell lysate (**Extended Data Fig. 6**). We subsequently analyzed SARS-CoV-2 sequencing data and RNA recombination events with “*ViReMa*” as described above [24, 25].

**Figure 5.**
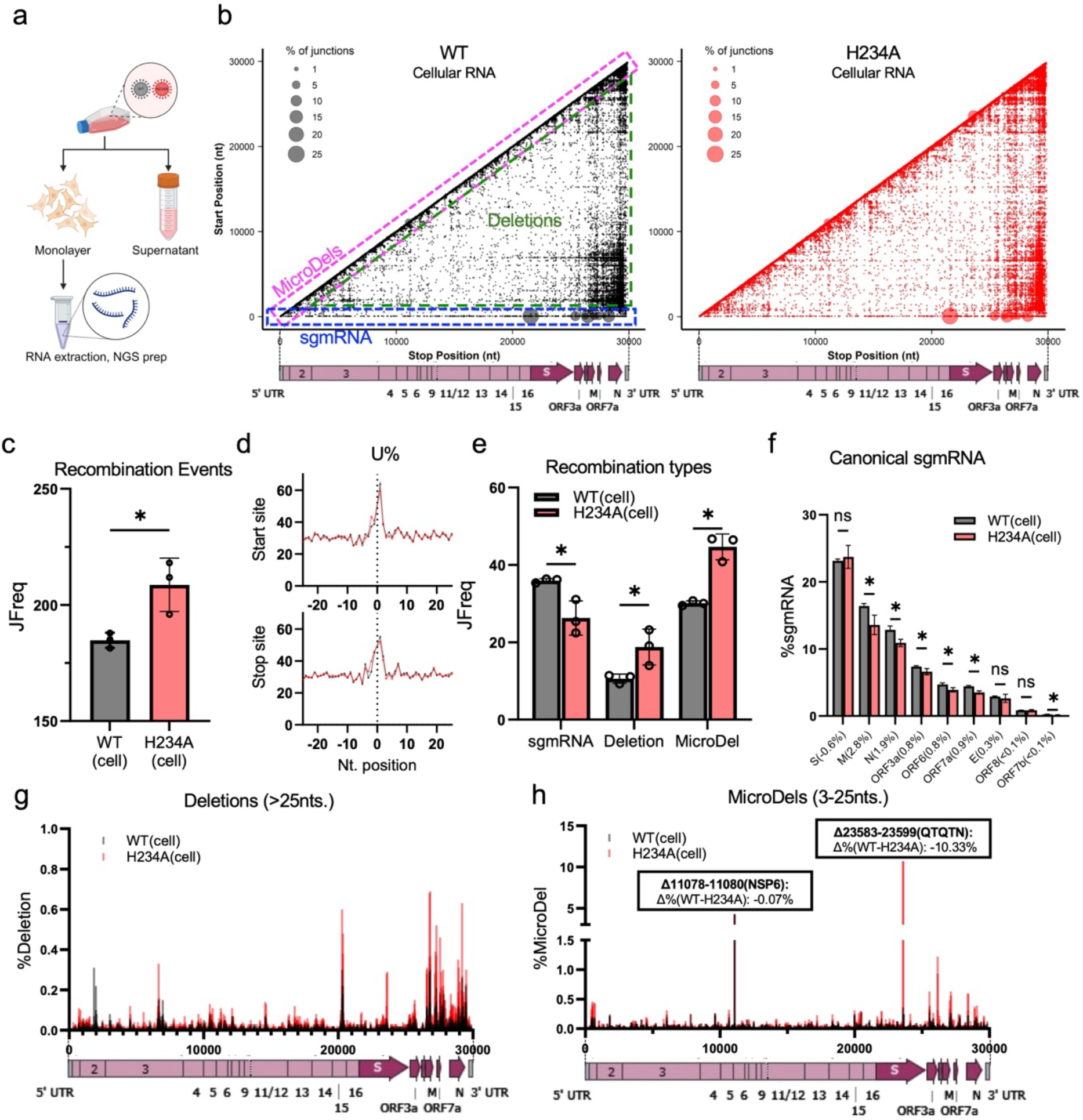
NSP15^H234A^ resulted in increased RNA recombination but reduced sgmRNA. (a) Schematic of RNA sequenced from monolayered cell infected by viruses. (b) 2D maps of recombination events and their frequencies from cell monolayer infected by WT (black) or H234A (red). Each event represents a recombination sequence mapped to a start position (Y-axis, donor site) and a stop position (X-axis, acceptor site). (c) Across the entire genome, H234A showed significantly higher recombination (JFreq: number of recombination reads per 104 viral genome reads) than WT. (d) H234A showed altered utilization frequencies of uridines flanking the recombination junctions. (e) H234A showed increased JFreq of deletion, micro-deletion (MicroDel), but reduced intracellular sgmRNA. (f) The most abundant 9 canonical sgmRNAs, their mean recombination rates and the percentage change of sgmRNA between WT and H234A; (g)1D map of deletion events (>25nts.) showed that H234A in general contained more genomic deletions than that of WT in cell lysate. (h)1D map of micro-deletions events (3-25nts.) showed H234A contained more genomic micro-deletions than WT in cell lysate. A substantial micro-deletion in NSP6 was shared for both WT and H234A. Another predominant microdeletion in spike protein is unique to H234A but not WT. Error bar: standard deviation. Two tailed T test with α = 0.05, N=3. *: P < 0.05, **: P < 0.01. ns : not significant.

Using two-dimensional scatter plots, we visualized the frequency and location of RNA recombination events relative to their start and stop position and normalized the frequency of each recombination event to sequencing depth at the junction (**Fig. 5b**). This depicts several types of recombination events including: 1) canonical and non-canonical sub-genomic mRNA, bound on the X axis and capturing events between the Transcription Regulatory Sequence (TRS)-leader and the rest of genome; (**Fig. 5b, blue box**) 2) micro-deletions (MicroDel) of <25 nucleotides along the X=Y axis (**Fig. 5b, magenta box**), 3) deletions (>25nts.) dispersed below the X=Y axis ((**Fig. 5b, green triangle**). In addition, we also detected other RNA recombination events such as end fusion (recombination between 3’-end of genome to 5’-start of genome) and insertion events (long and micro-insertions). However, no statistical differences were found between WT and H234A for these events, and they are therefore depicted separately for clarity (**Extended Data Fig. 7a**).

While RNA recombination was observed in both infection groups, NSP15^H234A^ infected cells produced significantly more RNA recombination than WT (**Fig. 5b**). Specifically, the NSP15^H234A^ showed a ∼16% increase in JFreq (recombination junctions per 10^4^ mapped viral reads) compared to WT (**Fig. 5c**). Having established that the lack of NSP15 activity increases RNA recombination, we next evaluated if the loss of NSP15 activity altered the uridine-enriched sequences flanking SARS-CoV-2 RNA recombination junctions (**Fig. 5d**). Following NSP15^H234A^ infection, we observed modest increase of uridine frequencies near RNA recombination start/stop sites mainly upstream of the junction, but no substantial differences in other nucleotides (**Extended Data Fig. 8**). These results suggest that endoU activity of NSP15 is not required for the uridine-favored RNA recombination in SARS-CoV-2.

Examination of the types of recombination indicated significant shifts between the WT and NSP15^H234A^ (**Fig. 5e**). For example, despite more overall abundant recombination events, NSP15^H234A^ infection produces less sub-genomic mRNAs as compared to WT (∼27% reduction). From the nine canonical sub-genomic mRNAs, we found that six had significant reduction compared to WT (**Fig. 5f**). We also use the linear viral sequencing across the intact TRS-L as a proxy for viral genomic RNA and evaluate the ratio of sub-genomic mRNA to viral genomic RNA (**Extended Data Fig. 9a**). While the frequency of intact TRS-L is similar between infections, NSP15^H234A^ showed a decreased TRS-L:TRS-B junction frequency that gives rise to sgmRNAs (**Extended Data Fig. 9b**). These changes result in a lower subgenomic/genomic RNA ratio and confirm reduced sgmRNA formation in NSP15^H234A^ compared to WT.

In contrast to the sgmRNA, the frequency of deletion (>25 nts.) and micro-deletions (<25nts.) was significantly higher in the NSP15^H234A^ as compared to WT (**Fig. 5e**), which are major contributors to defective viral genomes [14]. For deletions, the NSP15^H234A^ had a ∼77% increase in JFreq relative to WT. Examination of the deletion distribution found them spread throughout the genome for both infections, but more abundant in NSP15^H234A^ (**Fig. 5g**). Notably, despite more overall deletions in NSP15^H234A^, some sites were more abundant in WT cell lysate (e.g. start positions at nts.1854, 1988, and 6949). For micro-deletions, NSP15^H234A^ also showed an increase in JFreq (∼48%) compared to WT (**Fig. 5e**). Similar to deletions, NSP15^H234A^ micro-deletions showed increased frequencies and distribution, especially towards the 3’end of the viral genome (**Fig. 5h**). Together, this demonstrates that NSP15 plays a role in divergent regulation of RNA recombination events: limiting deletions and micro-deletions but also facilitating the formation of sgmRNAs.

Notably, while micro-deletion events were mainly with low frequency (<1.5%), two high frequency events were observed in our analysis. The first event, a high frequency micro-deletion between nts.11078-11080 (NSP6, ∼4.2%-4.3%), was shared by both WT and NSP15^H234A^. This micro-deletion recombination results in a single nucleotide change and a frame shift in the open reading frame. In contrast, the second high frequency micro-deletion recombination between nts. 23583-23599 was found to be ∼30-fold more frequent in NSP15^H234A^ than WT. This site corresponds to a deletion of the QTQTN motif found in the furin-cleavage loop of SARS-CoV-2 spike, a common attenuating mutation observed following passage in Vero cells [33]. We further scrutinized the accumulation of Δ11078-11080 (NSP6) and Δ23583-23599 (QTQTN) by sequencing the parental virus strains P1 stock (**Extended Data Fig. 10**). We found that both parental WT and NSP15^H234A^ had the Δ11078-11080 (NSP6) with comparable frequencies, which may be the result of T7/RDRP processibility in adjacent regions (11075-(U)_8_-11082) or a consistent sequencing artifact at this poly U region. In contrast, neither parental virus showed elevated frequency of Δ23583-23599 (QTQTN). Together, the loss of NSP15 activity led to the rapid accumulation of this micro-deletion conferring a fitness advantage in Vero cells.

### NSP15^H234A^ increased accumulation of defective viral genomes in virions

Cell associated RNA represents viral replication occurring in a complex intracellular environment under pressure by host anti-viral and immune processes [38]. In contrast, purified virions provide a controlled microenvironment to investigate if cellular RNA recombination events carry forward during infection, such as viral particles composed of defective viral genomes (DVGs) [42–44].To this end, we collected the supernatant from Vero E6 cells infected with WT SARS-CoV-2 or NSP15^H234A^, pelleted virus particles using sucrose cushion (**Fig. 6a**) and conducted NGS and bioinformatic analyses as described above.

**Figure 6.**
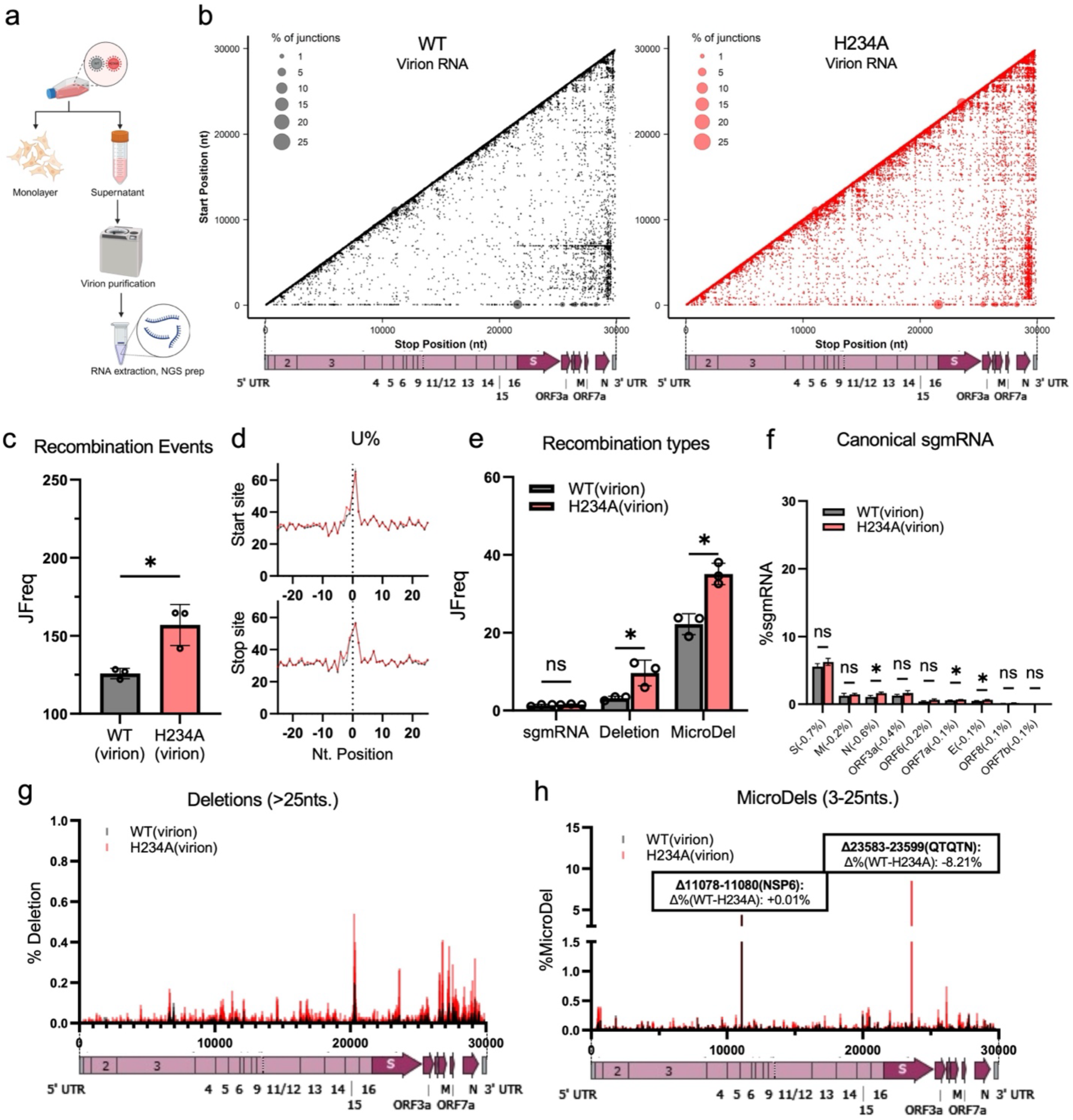
NSP15^H234A^ resulted in increased defective viral genomes (DVGs) in packaged virions. (a) Schematics of RNA sequenced from purified virions. (b) 2D maps of recombination events and their frequencies from virions of infected Vero E6. (c) The purified virions recapitulated the increased JFreq of H234A than that of WT. (d) H234A mutant virus particles recapitulated the altered recombination U frequencies similar to that of cell monolayer. (e) While little sgmRNA was encapsidated by virus particles, H234A particles contained more DVGs such as deletions and micro-deletions. (f) The most abundant 9 canonical sgmRNAs found in supernatant fraction. (g) 1D map of deletion events (>25nts.) showed that H234A in general contained more genomic deletions than that of WT in supernatant. (h) 1D map of micro-deletions events (3-25nts.) showed H234A contained more genomic micro-deletions than that of WT in supernatant. Error bar: standard deviation. Two tailed T test with α = 0.05, N=3. *: P < 0.05, **: P < 0.01. ns : not significant.

From purified virions, we found that viral RNA recombination trends recapitulated the events found in cell lysates (**Fig. 6b**). While the frequency of recombination events in virions decreased compared to cellular lysates, the NSP15^H234A^virions showed significantly increased (∼25%) JFreq compared to WT (**Fig. 6c**). Similarly, U-rich tracts were still the primary site for recombination in the virions with or without NSP15/endoU activity (**Fig. 6d, Extended Data Fig. 8**). These results confirm that the NSP15^H234A^ increased recombination without substantially compromising U-enriched tracts adjacent to recombination events in purified virions. Consistent with cellular data, no statistical differences were found between WT and NSP15^H234A^ for end-fusion or insertion events (**Extended Data Fig. 7b**).

Examination of the types of recombination revealed similarities and differences with cellular RNA analysis. Only trace amounts of sgmRNAs were detected in purified WT and NSP15^H234A^ virions (**Fig. 6e&f)**. This is consistent with sgmRNAs not being packaged into virion and reflects the relative purity of the virion preparation. Similar to viral cellular RNA, both deletions (>25nts.) and micro-deletions (<25nts.) were found to be significantly increased in the NSP15^H234A^ mutant relative to WT (∼200% and ∼58% increase of JFreq, respectively). Examining their distribution, the deletions had greater frequency and distribution across the genome in NSP15^H234A^ compared to the WT (**Fig. 6g**). Similarly, the micro-deletion rates were also more abundant and diverse in NSP15^H234A^ compared to WT (**Fig. 6h**). Notably, both highly abundant micro-deletions (Δ11078-11080 (NSP6) & Δ23583-23599 (QTQTN)) from cellular lysates were retained in the virions. In addition, the relative abundance was similar to cellular RNA findings with Δ11078-11080 at equivalent levels and Δ23583-23599 showed a ∼29-fold increase over WT in NSP15^H234A^. Importantly, the results demonstrate that the increased deletion and micro-deletions produced by NSP15^H234A^ infection can be recapitulated by virus packaging and carried forward as DVGs. These DVGs may augment immune responses during infections [14].

### NSP15^H234A^ reduced RNA recombination diversity and sgmRNA formation *in vivo*

Having demonstrated that the NSP15^H234A^ can significantly increase RNA recombination *in vitro*, we evaluated viral recombination events *in vivo*. Using total RNA from homogenized hamster lung tissue (**Fig. 7a**), we utilized tiled-clickseq and ViReMa to evaluate recombination events following WT and NSP15^H234A^ infection. For both WT and NSP15^H234A^ infected hamster lungs, we observed substantially fewer recombination events (**Fig. 7b-c**) as compared to cell lysate or virions (**Fig. 5-6**). This ∼2-3 fold reduction in recombination JFreq indicates that the *in vivo* environment restricts accumulation of recombination events. Importantly, reduced recombination frequency in hamsters is consistent with findings from human autopsy samples [14] and suggest more restrictive selection pressure impacts the accumulation of recombination events.

**Figure 7.**
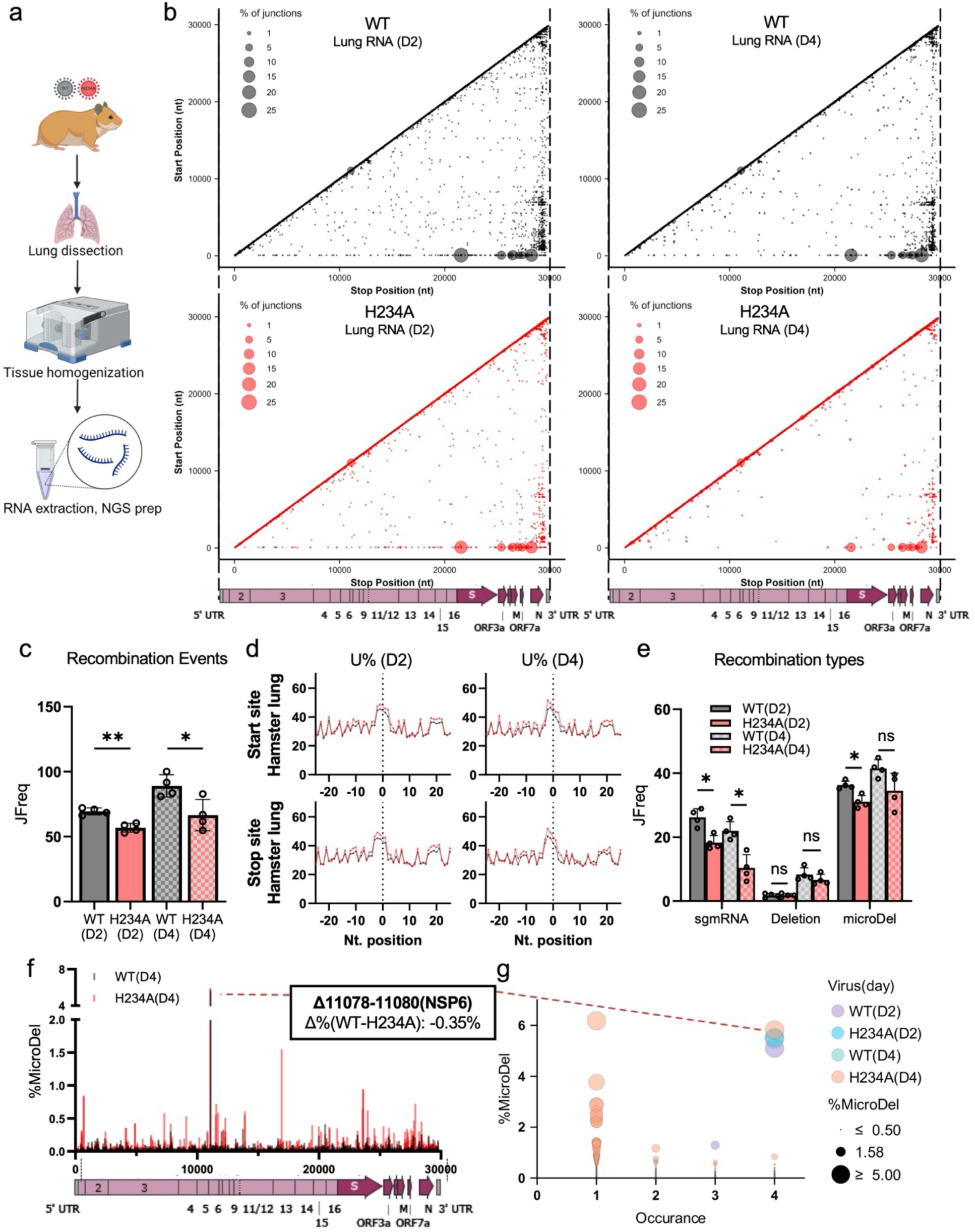
NSP15^H234A^ resulted in increased defective viral genomes (DVGs) in packaged virions. (a) Schematic of RNA sequenced from infected hamster lung. (b) 2D maps showing the coordinates of recombination events in hamster lung and the average frequency of each event at day 2 (D2) and day 4 (D4). (c) Across the entire genome, H234A showed slightly reduced recombination rate than correspondent WT control, at both D2 and D4; (d) H234A exhibited increased U utilization frequency flanking recombination junctions in hamster lungs. (e) H234A continued to significantly restrict the formation of sgmRNA in hamster lungs. H234A showed a trend of decreased rate of deletions and micro-deletions. (f) In hamster lungs, WT and H234A showed different patterns of microdeletions at D4: WT accumulated a myriad of near-background micro-deletions, while H234A contained fewer events but with higher frequencies. (g) Individual polymorphism of microdeletions in hamster lung tissue: H234A induced highly frequent micro-deletions that are specific to individual animal, whereas WT virus mainly contained the low frequency micro-deletions. Error bar: standard deviation. Two tailed T test with α = 0.05, N=4. *: P < 0.05, **: P < 0.01. ns : not significant.

In both day 2 and day 4 lung samples, the NSP15^H234A^ showed slightly reduced recombination events than WT, contrasting cellular and virion results. In the lung tissues, NSP15^H234A^ showed a modest increased in U% flanking RNA recombination start and stop sites relative to WT (**Fig. 7d, Extended Data Fig. 11**). This result is consistent with similar findings *in vitro* (**Extended Data Fig. 8**) and suggest that the endoU activity of NSP15 is not required for U-favored recombination of SARS-CoV-2.

Also consistent with *in vitro* findings, NSP15^H234A^ maintained significant reduction in sgmRNA (**Fig. 7e, Extended Data Fig. 12**). At both day 2 and 4, NSP15^H234A^ infected hamsters had reduced sgmRNA JFreq (30% and 52% respectively) relative to WT infected animals. Similarly, we observed a lowered ratio of sgmRNA to genomic RNA (**Extended Data Fig. 12c, d**) from the lungs of animals infected with NSP15^H234A^ relative to infection with WT. These results demonstrate that sgmRNA formation is reduced in the absence of NSP15 activity *in vivo*, similar to *in vitro* results.

Examining recombination types also found varying trends between *in vitro* and *in vivo* infection with WT and NSP15^H234A^. Both WT and NSP15^H234A^ had no significant differences between end-fusion, insertion and micro-insertion (**Extended Fig. 12a**), similar to cellular and virion analyses. However, both WT and NSP15^H234A^ demonstrate complexity in the context of deletion and micro-deletions. The NSP15^H234A^ infected hamster lungs exhibited a trend towards modest reductions in their total deletion and micro-deletions frequencies relative to WT (**Fig. 7e**). Examining further, the diversity and frequency of the recombination events highlight differences between WT and NSP15^H234A^. For example, at day 4, WT-infected hamster lungs accumulated one dominant (Δ11078-11080) and abundant low frequency micro-deletions (**Fig. 7f**). In contrast, the NSP15^H234A^ -infected lungs produced micro-deletions with less diversity, but with higher frequencies at specific sites (**Fig. 7f**). The high frequency, low diversity micro-deletions indicate strong selection as evidenced by the absence of Δ23583-23599 (ΔQTQTN) event in NSP15^H234A^ -infected lungs (Δ%wt-H234A=-0.1% on D2, -0.02% on D4). This ΔQTQTN mutation, highly penetrant in Vero cells, has been shown to be highly attenuated *in vivo* [33]. Similar to micro-deletions, the long deletion events in hamster lung followed the same trend (**Extended Data Fig. 12f,g**), with NSP15^H234A^ produced less abundant deletions but with higher frequencies at certain sites, especially at D4. We further demonstrate that NSP15^H234A^ infected animals have higher frequencies of individual deletion/micro-deletion, in spite of reduced diversity of events (**Extended Data Fig. 13**). Notably, the outcome of selection pressure varies across individual animals (**Fig. 7g**). Among four individual hamsters, NSP15^H234A^ -infected lungs gave rise to several high frequency micro-deletions that only occurred in a single animal. In contrast, WT-infected lungs contained background micro-deletions with low frequency in each individual, except for the common Δ11078-11080 (NSP6) event found in all animals from both groups at days 2 and 4. The increased frequency of deletion and micro-deletion events in NSP15^H234A^ infected lungs is consistent with *in vitro* findings, that the lack of NSP15 activity can still drive the accumulation of defective viral genomes *in vivo*. On the other hand, the reduced distribution of these events demonstrate that these defective viral genomes are shaped by a strong, individually divergent selective pressure *in vivo*.

## Discussions

While severity and lethality of COVID-19 have largely declined, SARS-CoV-2 remains a global health problem due to its ability to evade host immunity through evolution. Viral RNA recombination plays a role in this process, and we show here that SARS-CoV-2 recombines at a higher rate than other human CoVs (**Fig. 1b**). In addition, we confirm that hotspots for recombination occur at uridine-rich sites across all CoVs tested (**Fig. 1c**). Importantly, we also demonstrate a role for viral endonuclease NSP15 in balancing SARS-CoV-2 RNA recombination. In the absence of NSP15 catalytic activity (NSP15^H234A^), infected cells and purified virions accumulated more genomic deletions and micro-deletions. (**Fig. 5 & 6**). In contrast, the sgmRNA recombination is reduced in NSP15^H234A^ compared to WT. The *in vivo* model represents a stronger selective pressure and hence, lowered overall recombination rate (**Fig. 7**). *In vivo*, we recapitulate the reduced sgmRNA formation in NSP15^H234A^ compared to WT. In addition, the absence of NSP15 activity suppresses the diversity of recombination deletions but increases the frequency of a subset of events (**Fig. 7**). Combined with transcriptomic analyses, these results suggest that the loss of NSP15 activity renders a range of defective viral genomes (DVGs), which contribute to the observed increase in immune responses (**Fig. 4**). The induced antiviral state reduces viral replication, but also promotes immune-mediated damage, resulting in similar disease in hamsters infected with either mutant or WT virus (**Fig. 2&3**). Overall, our studies demonstrate that NSP15 catalytic activity plays a critical role in controlling host responses, facilitating sgmRNA formation, and limiting DVG accumulation during SARS-CoV-2 infection.

NSP15 endonuclease activity has previously been found to be a critical factor in controlling the host innate immune response [17, 18, 20–22]. Showing a preference for cleaving uridines and poly U tracts [27, 45–47], NSP15 endonuclease activity suppresses production of pathogen-associated molecular patterns and impairs immune sensing. The loss of NSP15 activity has also been associated with increased sensitivity to type I interferon treatment and attenuation of CoV replication [17, 18, 20–22]. Our study largely confirms these findings with enhanced sensitivity to type I IFN pretreatment (**Fig. 2**) and amplification of antiviral gene expression (**Fig. 4**) following NSP15^H234A^ infection. However, we also observed an increased expression of genes associated with inflammation and cytokines as well as significant lung pathology following NSP15^H234A^ infection. Together, our results show that the loss of NSP15 activity promotes an amplified immune response resulting in both viral suppression and immune mediated damage.

We predict that the amplified immune response observed in NSP15^H234A^ infection is the product of increased innate immune sensing and DVG production. Prior work with other CoVs has shown increases in dsRNA levels and other viral nucleic acids enhancing activation of immunes sensors [38, 48]. However, the production of DVGs in NSP15^H234A^ infection offers a mechanism that amplifies inflammation and damage responses. RNA viruses are known to produce DVGs which can shape the severity of disease [48]. Importantly, SARS-CoV-2 has already been shown to produce DVGs that promote host immune responses [14]. In this study, we find that loss of NSP15 activity increased deletions and micro-deletions significantly in viral RNA; importantly, these deletions and micro-deletions are observed both in viral RNA from cells and viral RNA packaged in virions. Acting as DVGs, NSP15^H234A^ infection produces amplified immune responses in terms of both antiviral activity and inflammation. This mechanism is consistent with our *in vivo* results finding reduced viral loads of NSP15^H234A^ despite significant weight loss, inflammation, and damage within the lung. Overall, our results indicate that NSP15 plays a critical role in limiting recombination and accumulation of immunogenic DVGs; the loss of the viral endoribonuclease activity augments both antiviral responses and immune mediated damage.

Our manuscript also provides critical insights into key elements of CoV recombination. SARS-CoV-2 has a higher rate of recombination than other human CoVs tested which may contribute to rapid development of novel variants (**Fig. 1b**). While more frequent RNA recombination may be unique to SARS-CoV-2, it is unclear if this trait is conserved in other sarbecoviruses and requires further study. Notably, for all CoVs tested, uridine rich tracts serve as the primary site for recombination (**Fig. 1c**) and the process operates independently of NSP15 activity. While we had initially postulated that NSP15 endoU serves to provide cleaved template RNA to facilitate recombination, we instead found accelerated recombination in its absence suggesting a key role in regulation of overall recombination.

Our results also provide experimental insights to connect NSP15 endoU activity to the correct formation of SARS-CoV-2 canonical sgmRNAs. The coronavirus transcription regulatory sequence (TRS) is a conserved RNA motif that resides at stem loop 3 of virus 5’UTR [49]. The production of subgenomic mRNA (sgmRNA) relies on the correct recognition of the complementarity between leader TRS (TRS-L) and the recombination to body TRS (TRS-B) [50, 51]. The conserved Sarbecoviruses “AACGAAC” TRS-L motif [52] is A-rich in positive sense and U-rich in negative sense viral RNAs, which is also flanked by A/U-rich sequences both up- and down-stream. It is conceivable that the endoU activity of NSP15 may play a role in mediating the template-switching between +gRNA and -sgmRNA to give rise to +sgmRNA. Indeed, previous studies speculated that NSP15 cleavage of TRS is required to form sgmRNA [53, 54]. Our data demonstrated that the loss of endoU activity significantly down-regulated the formation of TRS-L and TRS-B recombination both *in vivo* (**Fig.5**) and *in vitro* (**Fig. 7**). This provides experimental evidence that NSP15 plays a critical role to “proofread” the correct recombination between TRS-L and TRS-B.

Our results also provide mechanistic insight into the role of NSP15 beyond viral IFN antagonist. Our data show augmented host immune responses in the NSP15^H234A^, but also a shift in DVG formation. The dual impact of limiting innate immune sensing and production of immunogenic DVGs highlight the crucial role for NSP15 during SARS-CoV-2 infection. It is also possible that NSP15 activity varies across the CoV family, modulating the levels of recombination and possibly explaining the increased recombination observed in SARS-CoV-2. Notably, it also questions the safety of targeting NSP15 for drug and therapeutic treatment. While targeting NSP15 activity may attenuate viral replication, it may also promote immune mediated damage as a byproduct of treatment. In addition, NSP15 targeting may increase viral recombination permitting more rapid formation of resistance to this and other treatments. Given the mutagenesis concerns associated with molnupiravir treatment, similar safety challenges might be associated with NSP15-targeted treatments.

Overall, this research provides a detailed exploration of SARS-CoV-2 recombination *in vitro* and *in vivo*. Our results confirm higher recombination in SARS-CoV-2 primarily at uridine-rich tracts. Importantly, we show that SARS-CoV-2 NSP15 endonuclease activity is key to balancing recombination in cells, in virions, and *in vivo*. Together, the work highlights novel elements of CoV recombination and novel mechanistic insights into how NSP15 modulates host immunity and defective viral genome production.

## Methods

### Cell culture

Vero E6 cells were cultured in high glucose DMEM (Gibco) supplemented with 10% fetal bovine serum (HyClone) and 1x antibiotic-antimycotic (Gibco). Calu3 2B4 cells were cultured in high glucose DMEM supplemented with 10% defined fetal bovine serum, 1mM sodium pyruvate (Gibco), and 1x antibiotic-antimycotic. Cells were maintained at 37°C in a humidified incubator with 5% CO_2_.

### Viruses

The recombinant and mutant SARS-CoV-2 viruses were generated based on the USA-WA1/2020 sequence provided by the World Reference Center for Emerging Viruses and Arboviruses, which was originally obtained by the Center for Disease Control and Prevention [55]. The Nsp15 mutant was constructed with restriction enzyme-based cloning techniques and our reverse genetic system as previously described [56]. Virus stocks were amplified using Vero E6 cells. Viral RNA was extracted from recovered viruses, and the mutation was verified using next generation sequencing as previously described [57].

### *In vitro* infection

In vitro infection of Vero E6 and Calu3 2B4 cells was performed as previously described [33]. Briefly, Vero E6 or Calu3 2B4 cells were seeded in a 6-well plate format. For experiments involving IFN-I pre-treatment, Vero E6 cells were treated with 100 units of Universal Type I IFN for 16 hours prior to infection. Cell growth media was removed and infected with either WT or mutant SARS-CoV-2 virus at an MOI of 0.01 for 45 min at 37°C.

Following adsorption, cells were washed three times with phosphate buffered saline, and fresh growth medium was added. Three or more biological replicates were collected at each time point. Viral titers of the samples were subsequently determined using focus forming assay as previously described [32, 58].

Following *in vitro* infection, culture supernatant was harvested, clarified, and virus particles were pelleted by ultracentrifugation with previously established methods [33].

### *In vivo* infection

Three-to-four-week-old male golden Syrian hamsters were purchased from Envigo. Animals were housed in ventilated cages prior to the study. Animals were intranasally infected with 10^5^ FFU of WT or H234A in 100-ul inoculum or mock-infected with PBS. Animals were monitored daily for weight loss and signs of clinical disease for up to seven days post infection (DPI). On days 2, 4, and 7, five animals from each group were anesthetized with isoflurane and nasal washed with PBS and subsequently euthanized with CO_2_ for organ collection. Lung lobes were collected in either PBS for viral titers, RNAlater (Invitrogen #AM7021) for NGS/gene expression, or 10% phosphate-buffered formalin (Fisher #SF100) for histopathology.

### Histology

Left lung lobes were collected and fixed in 10% buffered formalin for at least 7 days. Fixed tissues were paraffin-embedded, sections cut into 5-µM thickness and stained with hematoxylin and eosin on a SAKURA VIP 6 tissue processor at the University of Texas Medical Branch Surgical Pathology Laboratory. For viral antigen staining, tissue sections were deparaffinized and reacted with SARS-CoV-2 N-specific primary antibody and incubated with a secondary HRP-conjugated anti-rabbit antibody as previously described [58]. Viral antigen was visualized and scored blinded on a scale of 0 (none) to 3 (most) in 0.25 increments with scores averaged from at least two sections from each hamster.

### Virus quantitation

For in vitro samples, viral titers were measured using focus forming assay as previously described [58]. Briefly, hamster lung lobes were homogenized with zirconia beads in a MagNA Lyser instrument (Roche Life Science) and clarified with low-speed centrifugation. Vero E6 cells were seeded in 96-well plates to achieve 100% confluency at the time of titration. A 10-fold serial dilution was performed for virus-containing supernatant, and 20 uL of the dilutions were transferred to Vero E6 cells after the culture medium was removed. Cells were incubated for 45 min at 37 °C with 5% CO2 to allow adsorption before 0.85% methylcellulose overlay was added. After removing the overlay, cells were washed 3 times with PBS before fixation in 10% buffered formalin for 30 min at room temperature. Cells were permeabilized and incubated with SARS-CoV-2 nucleocapsid antibody (Cell Signaling) followed by Alexa FluorTM 555-conjugated α-mouse secondary antibody (Invitrogen). Fluorescent foci images were captured on Cytation 7 imaging multi-mode reader (BioTek) and foci were counted with ImageJ.

### Next Generation Sequencing (NGS) libraries

For NGS analyses, RNA template was extracted from infected cells, supernatant or animal tissue with Direct-zol RNA miniprep kits (Zymo Research).

To sequence different human coronaviruses, a random hexamer (N_6_) primer was used with standard “ClickSeq” [23]. In brief, viral RNAs were extracted from cell lysate and reverse transcribed with N_6_ and 1:35 azido-ddNTP:dNTP ratio. The azido-ddNTP-terminated cDNAs were “click-ligated” with a 3’-alkyne modified adapter. The ligated cDNAs then underwent final PCR to fulfill illumina library structure. Gel selected libraries were single-end sequenced with illumina NextSeq 2K.

To investigate the recombination rate of SARS-CoV-2, we used “Tiled-ClickSeq” [26] approach, which uses >300 primers specific to SARS-CoV-2 genome and ClickSeq components [23, 59] to achieve sensitive detection and even coverage of coronavirus genome. The extracted RNAs from cell or animal tissue were used as template for Tiled-ClickSeq libraries with standard protocol [26].

To understand the transcriptomic changes of infected animals, the extracted total cellular RNA from hamster lungs were subjected to PolyA-ClickSeq [39], which utilizes a semi-anchored oligo(dT) primer (5’-(T)_21_-3’) to specifically target polyA tails of cellular mRNA and a 1:5 azido-ddNTP:dNTP ratio to ensure sufficient termination of cDNA. The PolyA-ClickSeq library was constructed with previously established protocol [60] and gel selected libraries were single-end sequenced with ElementBio Aviti.

### Bioinformatics

Raw reads sequenced from Tiled-ClickSeq libraries are processed and analyzed with previously established bioinformatic pipelines *TCS* (https://github.com/andrewrouth/TCS) with parameters *“-p PMV*”. In brief, after initial quality filter and trimming of illumina adapter, the detected primer sequences from R2 reads will subsequently be trimmed from respective R1 reads (therefore excluding potential primer-genome recombination). The processed R1 reads are then mapped to SARS-CoV-2 reference genome (NCBI reference: NC_045512.2). The same processed R1 reads are also analyzed with ViReMa (version 0.28) (https://sourceforge.net/projects/virema/) with parameters “--*ErrorDensity 2,30 –Seed 25 --Defuzz --X 3”.* The ViReMa-output BED files are further analyzed to categorize different recombination events with custom python script “*plot_cs_freq.py*” (included in the *TCS* package), which defines the length of microindel to be within 25nts. 2D-maps of ViReMa-mapped recombination events are plotted with *ViReMaShiny* [61] (https://routhlab.shinyapps.io/ViReMaShiny/) with modifications for cosmetic and style.

Raw reads sequenced from random primed libraries are processed with *fastp* [62] for adapter removal and quality control. To remove potential artifacts of random hexamer, all R1 reads were trimmed by 8 bases from 3’-direction (“*-t 8*”). For each virus, the corresponding virus genome was first polished with Pilon[63] with sequenced reads to improve mapping efficiency. This is followed by *ViReMa* mapping of recombination reads to the *Pilon*[63]-polished viral genomes (NCBI reference number SARS-CoV-2: NC_045512.2; MERS-CoV: NC_019843.3; hCoV-229E: NC_002645.1; hCoV-OC43: ATCC VR-1558) with the same parameters stated above.

Raw reads sequenced from PolyA-ClickSeq libraries are processed and analyzed with previously established *DPAC* [40] pipeline (https://github.com/andrewrouth/DPAC/) with parameters “*-p PMBCD*”, which detects and processes the polyA-containing reads and maps the upstream sequence of polyA tail to host reference. In this study, a PolyA-site clustering data base was curated based on the published reference genome and annotation of *Mesocricetus auratus* (Genbank accession number: GCA_000349665.1). Due to the incompleteness of the reference genome, the genome mapping criterion was slightly loosened in hisat2 [64] stage with parameter *“--score-min L,0,-0.3*”. Differential gene expression profiles of mock (1X PBS), wt, and H234A infected hamster lung tissues were then analyzed with DESeq2 [65] that has been integrated in the *DPAC* pipeline (D stage) to reveal the normalized read count of each annotated gene.

For differential gene expression analyses, hierarchical gene clustering was conducted with *Cluster 3.0* (http://bonsai.hgc.jp/~mdehoon/software/cluster/). This is followed by TreeView (http://jtreeview.sourceforge.net/) to visualize the heat map and gene clusters.

For the significant (*p*_adj_ < 0.1, |log2FC| > 1) differentially expressed genes of wt(D2) vs. H234A(D2), gene ontology assay was conducted with DAVID [41] (https://david.ncifcrf.gov/home.jsp) to highlight the most direct GO terms in biological process, cellular component, and molecular function. Pathway discovery and illustration were conducted with KEGG Pathway Database (https://www.genome.jp/kegg/pathway.html).

### Data Availability

The raw sequencing data of this study are available in the NCBI sequence read archive (SRA) with accession number: **PRJNA1131338, PRJNA1154272**. MERS-CoV data are reanalyzed from existing SRA project: **PRJNA623016**[15]. Raw reads count and mapping rate are listed in **Supplementary Table S1**.

## Acknowledgements

Special thanks to UTMB next generation sequencing core staff (Haiping Hao, Jill K. Thompson) for sequencing support.

## Funding

Research was supported by grants from NIAID of the NIH to (AI168232, AI153602, and U19AI171413) to VDM. ZY and ALR was supported by an Institute of Human Infection and Immunity at UTMB COVID-19 Research Fund. Research was also supported by Burroughs Wellcome Fund (Investigators in Pathogenesis of Infectious Disease) to VDM and Chan Zuckerberg Initiative (Single-Cell Analysis Inflammation Grant) to VDM & ALR.

## Competing Interests

A.L.R. is a co-founder and co-owner of “ClickSeq Technologies LLC.”, a next-generation sequencing service provider of ClickSeq protocols and downstream analyses such as those presented in this manuscript. VDM have filed a patent on the reverse genetic system for SARS-CoV-2. All other authors declare no conflicts of interest.

## Author contributions

Conceptualization: YZ, YPA, VDM, ALR

Formal analysis: YZ, YPA, KGL, REA, DHW, BAJ, ALR, VDM

Funding acquisition: ALR, VDM

Investigation: YZ, YPA, KGL, REA, LKE, WMM, AMM, ALM, JTM, DHW, BAJ

Methodology: YZ, YPA, KGL, REA, DHW, ALR, VDM

Project Administration: ALR, VDM

Supervision: BAJ, ALR, VDM

Visualization: YZ, YPA, ALR, VDM

Writing – original draft: YZ, YPA, VDM

Writing – review and editing: YZ, YPA, KGL, REA, DHW, BAJ, ALR, VDM

## References

1. WHO COVID-19 dashboard. 2024 [cited 2024 7/19]; Available from: https://data.who.int/dashboards/covid19/cases.

2. Gralinski, L.E. and V.D. Menachery, Return of the Coronavirus: 2019-nCoV. Viruses, 2020. 12(2).

3. Amanat, F. and F. Krammer, SARS-CoV-2 Vaccines: Status Report. Immunity, 2020. 52(4): p. 583–589.

4. DeGrace, M.M., et al., Defining the risk of SARS-CoV-2 variants on immune protection. Nature, 2022. 605(7911): p. 640–652.

5. Plante, J.A., et al., The variant gambit: COVID-19’s next move. Cell Host Microbe, 2021. 29(4): p. 508–515.

6. Davis, H.E., et al., Author Correction: Long COVID: major findings, mechanisms and recommendations. Nat Rev Microbiol, 2023. 21(6): p. 408.

7. Bowe, B., Y. Xie, and Z. Al-Aly, Acute and postacute sequelae associated with SARS-CoV-2 reinfection. Nat Med, 2022. 28(11): p. 2398–2405.

8. Fehr, A.R. and S. Perlman, Coronaviruses: an overview of their replication and pathogenesis. Methods Mol Biol, 2015. 1282: p. 1–23.

9. Minskaia, E., et al., Discovery of an RNA virus 3’->5’ exoribonuclease that is critically involved in coronavirus RNA synthesis. Proc Natl Acad Sci U S A, 2006. 103(13): p. 5108–13.

10. Wells, H.L., et al., The coronavirus recombination pathway. Cell Host Microbe, 2023. 31(6): p. 874–889.

11. Sola, I., et al., Continuous and Discontinuous RNA Synthesis in Coronaviruses. Annu Rev Virol, 2015. 2(1): p. 265–88.

12. Keck, J.G., et al., In vivo RNA-RNA recombination of coronavirus in mouse brain. J Virol, 1988. 62(5): p. 1810–3.

13. Makino, S., et al., High-frequency RNA recombination of murine coronaviruses. J Virol, 1986. 57(3): p. 729–37.

14. Zhou, T., et al., Generation and Functional Analysis of Defective Viral Genomes during SARS-CoV-2 Infection. mBio, 2023. 14(3): p. e0025023.

15. Gribble, J., et al., The coronavirus proofreading exoribonuclease mediates extensive viral recombination. PLoS Pathog, 2021. 17(1): p. e1009226.

16. Deng, X., et al., Coronavirus nonstructural protein 15 mediates evasion of dsRNA sensors and limits apoptosis in macrophages. Proc Natl Acad Sci U S A, 2017. 114(21): p. E4251–E4260.

17. Hackbart, M., X. Deng, and S.C. Baker, Coronavirus endoribonuclease targets viral polyuridine sequences to evade activating host sensors. Proc Natl Acad Sci U S A, 2020. 117(14): p. 8094–8103.

18. Deng, X., et al., Coronavirus Endoribonuclease Activity in Porcine Epidemic Diarrhea Virus Suppresses Type I and Type III Interferon Responses. J Virol, 2019. 93(8).

19. Otter, C.J., et al., SARS-CoV-2 nsp15 endoribonuclease antagonizes dsRNA-induced antiviral signaling. bioRxiv, 2023.

20. Kindler, E., et al., Early endonuclease-mediated evasion of RNA sensing ensures eZicient coronavirus replication. PLoS Pathog, 2017. 13(2): p. e1006195.

21. Volk, A., et al., Coronavirus Endoribonuclease and Deubiquitinating Interferon Antagonists DiZerentially Modulate the Host Response during Replication in Macrophages. J Virol, 2020. 94(11).

22. Otter, C.J., et al., SARS-CoV-2 nsp15 endoribonuclease antagonizes dsRNA-induced antiviral signaling. Proceedings of the National Academy of Sciences, 2024. 121(15): p. e2320194121.

23. Routh, A., et al., ClickSeq: Fragmentation-Free Next-Generation Sequencing via Click Ligation of Adaptors to Stochastically Terminated 3’-Azido cDNAs. Journal of molecular biology, 2015. 427(16): p. 2610–2616.

24. Sotchef, S., et al., ViReMa: a virus recombination mapper of next-generation sequencing data characterizes diverse recombinant viral nucleic acids. Gigascience, 2023. 12.

25. Routh, A. and J.E. Johnson, Discovery of functional genomic motifs in viruses with ViReMa-a Virus Recombination Mapper-for analysis of next-generation sequencing data. Nucleic Acids Res, 2014. 42(2): p. e11.

26. Jaworski, E., et al., Tiled-ClickSeq for targeted sequencing of complete coronavirus genomes with simultaneous capture of RNA recombination and minority variants. Elife, 2021. 10: p. e68479.

27. Pillon, M.C., et al., Cryo-EM structures of the SARS-CoV-2 endoribonuclease Nsp15 reveal insight into nuclease specificity and dynamics. Nat Commun, 2021. 12(1): p. 636.

28. Comar, C.E., et al., MERS-CoV endoribonuclease and accessory proteins jointly evade host innate immunity during infection of lung and nasal epithelial cells. Proc Natl Acad Sci U S A, 2022. 119(21): p. e2123208119.

29. Xie, X., et al., Engineering SARS-CoV-2 using a reverse genetic system. Nat Protoc, 2021. 16(3): p. 1761–1784.

30. Xie, X., et al., An Infectious cDNA Clone of SARS-CoV-2. Cell Host Microbe, 2020. 27(5): p. 841–848 e3.

31. Lokugamage, K.G., et al., Type I Interferon Susceptibility Distinguishes SARS-CoV-2 from SARS-CoV. J Virol, 2020. 94(23).

32. Johnson, B.A., et al., Nucleocapsid mutations in SARS-CoV-2 augment replication and pathogenesis. PLoS Pathog, 2022. 18(6): p. e1010627.

33. Vu, M.N., et al., QTQTN motif upstream of the furin-cleavage site plays a key role in SARS-CoV-2 infection and pathogenesis. Proc Natl Acad Sci U S A, 2022. 119(32): p. e2205690119.

34. Garvanska, D.H., et al., The NSP3 protein of SARS-CoV-2 binds fragile X mental retardation proteins to disrupt UBAP2L interactions. EMBO Rep, 2024. 25(2): p. 902–926.

35. Schindewolf, C., et al., SARS-CoV-2 Uses Nonstructural Protein 16 To Evade Restriction by IFIT1 and IFIT3. J Virol, 2023. 97(2): p. e0153222.

36. Deng, X., et al., Inactivating Three Interferon Antagonists Attenuates Pathogenesis of an Enteric Coronavirus. J Virol, 2020. 94(17).

37. Deng, X. and S.C. Baker, An “Old” protein with a new story: Coronavirus endoribonuclease is important for evading host antiviral defenses. Virology, 2018. 517: p. 157–163.

38. Otter, C.J., et al., SARS-CoV-2 nsp15 endoribonuclease antagonizes dsRNA-induced antiviral signaling. Proc Natl Acad Sci U S A, 2024. 121(15): p. e2320194121.

39. Routh, A., et al., Poly(A)-ClickSeq: click-chemistry for next-generation 3-end sequencing without RNA enrichment or fragmentation. Nucleic Acids Res, 2017. 45(12): p. e112.

40. Routh, A., DPAC: A Tool for DiZerential Poly(A)-Cluster Usage from Poly(A)-Targeted RNAseq Data. G3 (Bethesda), 2019. 9(6): p. 1825–1830.

41. Sherman, B.T., et al., DAVID: a web server for functional enrichment analysis and functional annotation of gene lists (2021 update). Nucleic Acids Res, 2022. 50(W1): p. W216–W221.

42. Zhou, Y. and A. Routh, Mapping RNA-capsid interactions and RNA secondary structure within virus particles using next-generation sequencing. Nucleic Acids Res, 2020. 48(2): p. e12.

43. Vignuzzi, M. and C.B. López, Defective viral genomes are key drivers of the virus– host interaction. Nature Microbiology, 2019. 4(7): p. 1075–1087.

44. Lopez, C.B., Defective viral genomes: critical danger signals of viral infections. J Virol, 2014. 88(16): p. 8720–3.

45. Frazier, M.N., et al., Flipped over U: structural basis for dsRNA cleavage by the SARS-CoV-2 endoribonuclease. Nucleic Acids Res, 2022. 50(14): p. 8290–8301.

46. Frazier, M.N., et al., Characterization of SARS2 Nsp15 nuclease activity reveals it’s mad about U. Nucleic Acids Res, 2021. 49(17): p. 10136–10149.

47. Bhardwaj, K., L. Guarino, and C.C. Kao, The severe acute respiratory syndrome coronavirus Nsp15 protein is an endoribonuclease that prefers manganese as a cofactor. J Virol, 2004. 78(22): p. 12218–24.

48. Wang, X. and B. Zhu, SARS-CoV-2 nsp15 preferentially degrades AU-rich dsRNA via its dsRNA nickase activity. Nucleic Acids Res, 2024. 52(9): p. 5257–5272.

49. Miao, Z., et al., Secondary structure of the SARS-CoV-2 5’-UTR. RNA Biol, 2021. 18(4): p. 447–456.

50. Malone, B., et al., Structures and functions of coronavirus replication-transcription complexes and their relevance for SARS-CoV-2 drug design. Nat Rev Mol Cell Biol, 2022. 23(1): p. 21–39.

51. Hartenian, E., et al., The molecular virology of coronaviruses. J Biol Chem, 2020. 295(37): p. 12910–12934.

52. Yang, Y., et al., Characterizing Transcriptional Regulatory Sequences in Coronaviruses and Their Role in Recombination. Mol Biol Evol, 2021. 38(4): p. 1241–1248.

53. Liang, J., et al., How the Replication and Transcription Complex Functions in Jumping Transcription of SARS-CoV-2. Front Genet, 2022. 13: p. 904513.

54. Li, X., et al., A Negative Feedback Model to Explain Regulation of SARS-CoV-2 Replication and Transcription. Front Genet, 2021. 12: p. 641445.

55. Harcourt, J., et al., Severe acute respiratory syndrome coronavirus 2 from patient with coronavirus disease, United States. Emerging infectious diseases, 2020. 26(6): p. 1266.

56. Xie, X., et al., Engineering SARS-CoV-2 using a reverse genetic system. Nature Protocols, 2021. 16(3): p. 1761–1784.

57. Jaworski, E., et al., Tiled-ClickSeq for targeted sequencing of complete coronavirus genomes with simultaneous capture of RNA recombination and minority variants. Elife, 2021. 10.

58. Garvanska, D.H., et al., The NSP3 protein of SARS-CoV-2 binds fragile X mental retardation proteins to disrupt UBAP2L interactions. EMBO reports, 2024. 25(2): p. 902–926.

59. Jaworski, E. and A. Routh, ClickSeq: Replacing Fragmentation and Enzymatic Ligation with Click-Chemistry to Prevent Sequence Chimeras, in Next Generation Sequencing: Methods and Protocols, S.R. Head, P. Ordoukhanian, and D.R. Salomon, Editors. 2018, Springer New York: New York, NY. p. 71–85.

60. Elrod, N.D., et al., Development of Poly(A)-ClickSeq as a tool enabling simultaneous genome-wide poly(A)-site identification and diZerential expression analysis. Methods, 2019. 155: p. 20–29.

61. Yeung, J. and A.L. Routh, ViReMaShiny: an interactive application for analysis of viral recombination data. Bioinformatics, 2022. 38(18): p. 4420–4422.

62. Chen, S., et al., fastp: an ultra-fast all-in-one FASTQ preprocessor. Bioinformatics, 2018. 34(17): p. i884–i890.

63. Walker, B.J., et al., Pilon: an integrated tool for comprehensive microbial variant detection and genome assembly improvement. PLoS One, 2014. 9(11): p. e112963.

64. Kim, D., et al., Graph-based genome alignment and genotyping with HISAT2 and HISAT-genotype. Nat Biotechnol, 2019. 37(8): p. 907–915.

65. Love, M.I., W. Huber, and S. Anders, Moderated estimation of fold change and dispersion for RNA-seq data with DESeq2. Genome Biol, 2014. 15(12): p. 550.

