## Supplemental Data for "SARS-CoV-2 EndoU-ribonuclease regulates RNA recombination and impacts viral fitness"

Extended Data Figures 1-13.

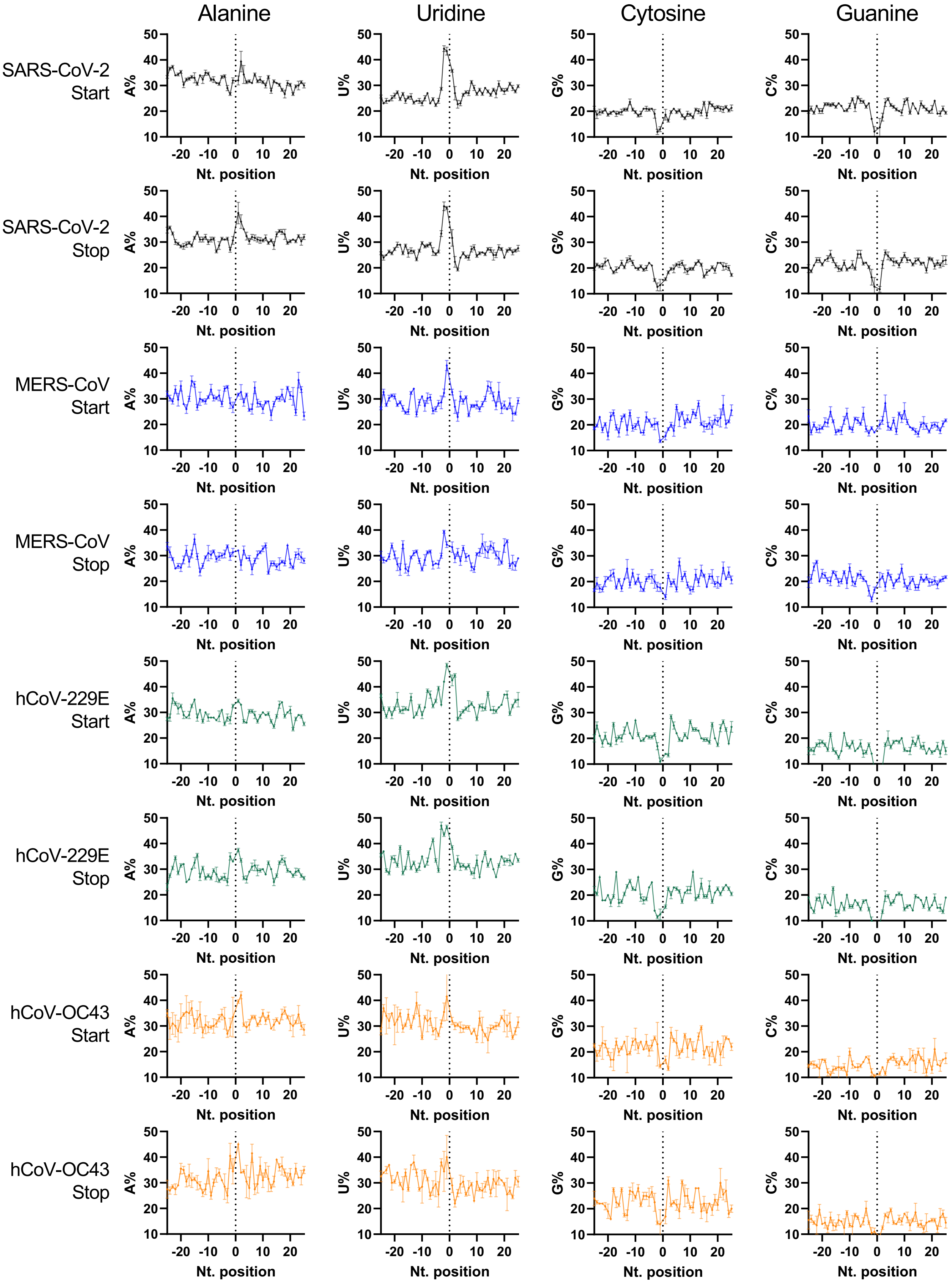


**Extended Data Figure 1. The frequency of A, U, G, C at each nucleotide position flanking the recombination start (donor) and stop (acceptor) sites.** Nucleotide frequency calculations were weighted by the abundance (read number) of each recombination event. sgmRNA events were excluded from this calculation due to their predominant abundance and putatively different mechanisms from other recombination events. All four human coronaviruses showed favorability of U-rich sequences flanking recombination junctions.  Error bar: standard deviation. N=3 for SARS-CoV-2 and MERS-CoV; N=2 for hCoV-229E and hCoV-OC43

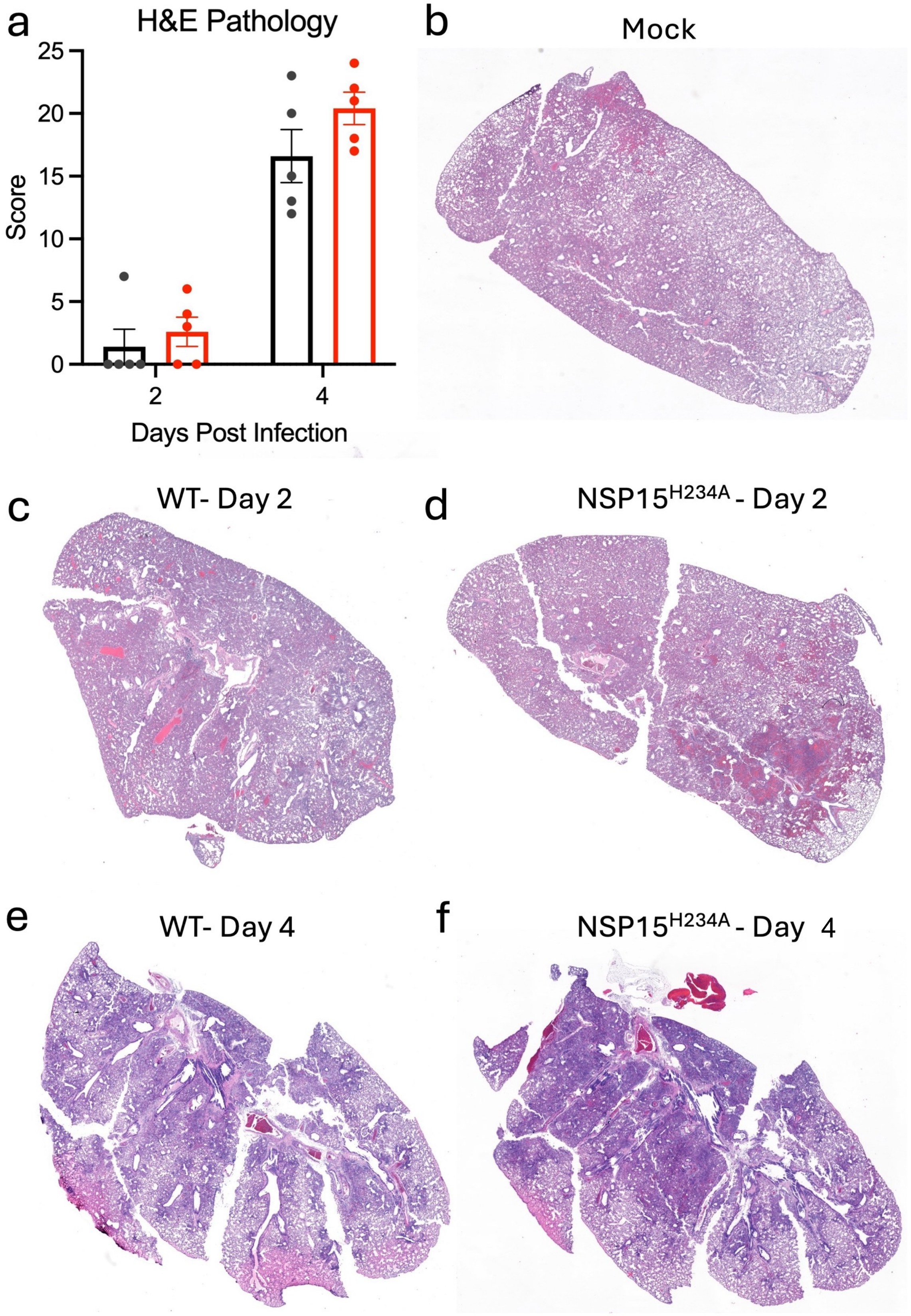


**Extended Data Figure 2. NSP15$^{H234A}$ has increased infiltration and immune damage than control.** Hamster lungs infected with WT or NSP15$^{H234A}$ were sectioned and stained (H&E) and evaluated for disease in damage by a board certified pathogist. (a) Two lung sections per animal per condition (n=5) were scored for pathology and infiltration. (b-f) Representative lung sections from (b) mock, day 2 and 4 infected with WT (c, e) and NSP15$^{H234A}$ (d, f).

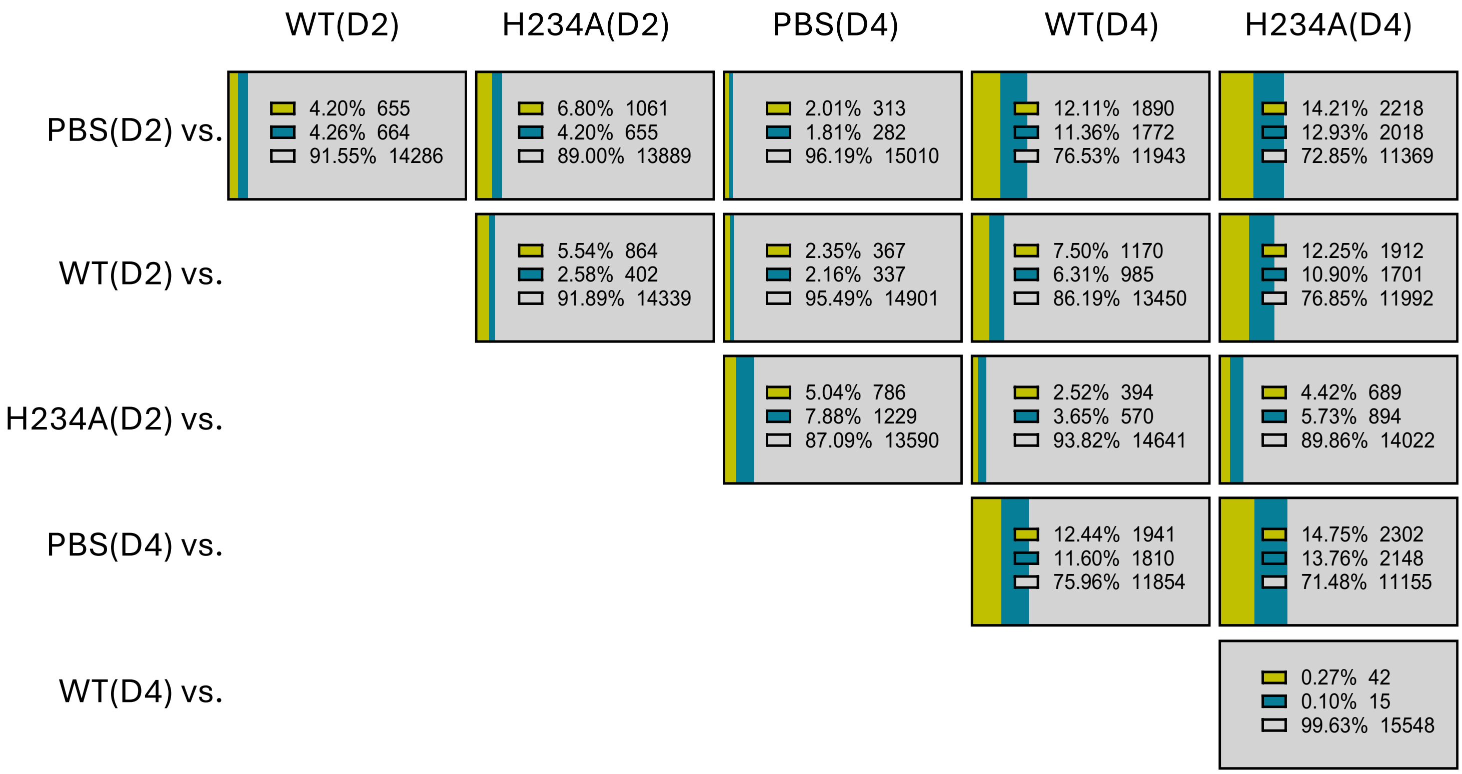


**Extended Data Figure 3**: Differentially expressed genes ($P_{adj}$< 0.1, |log2FC| > 0.585, N=3) between groups. Day 2 (D2) marks the largest difference between WT and H234A.

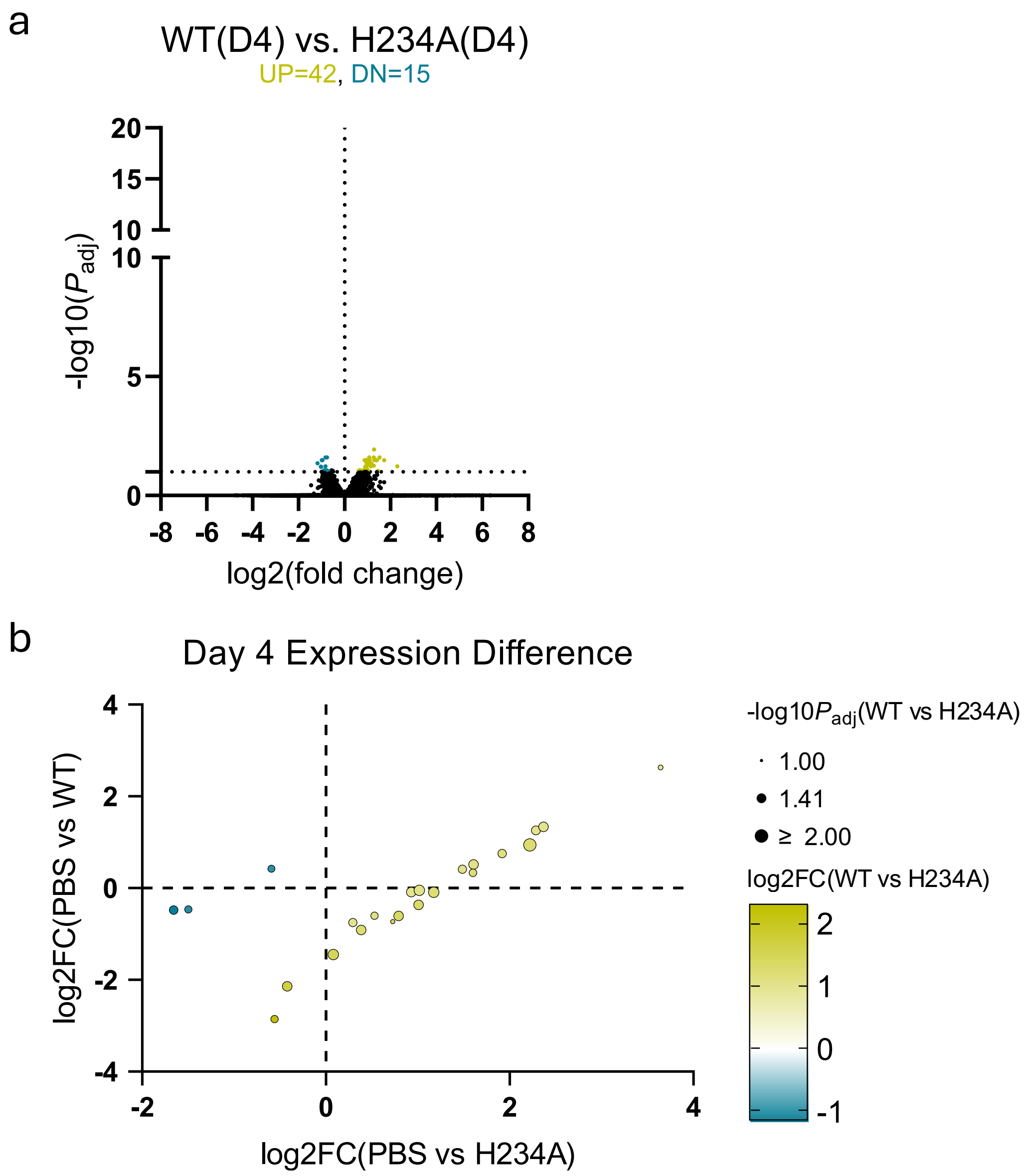


**Extended Data Figure 4.** (a) differentially expressed genes of WT vs H234A (D4). (b) Correlation of differentially expressed genes between WT and H234A at Day 4.

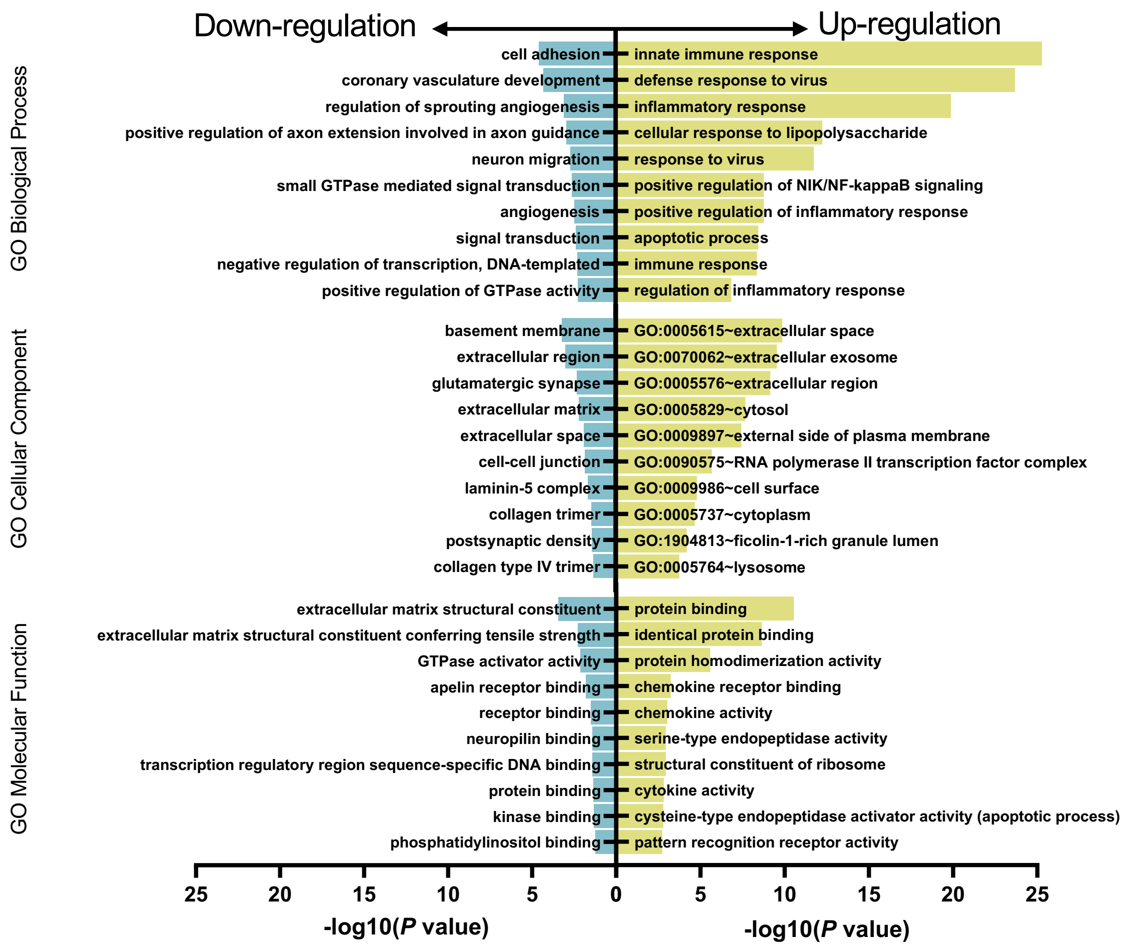


**Extended Data Figure 5.** Top 10 GO terms (by *P*-value) of up- or down-regulated genes of WT vs. H234A at D2.

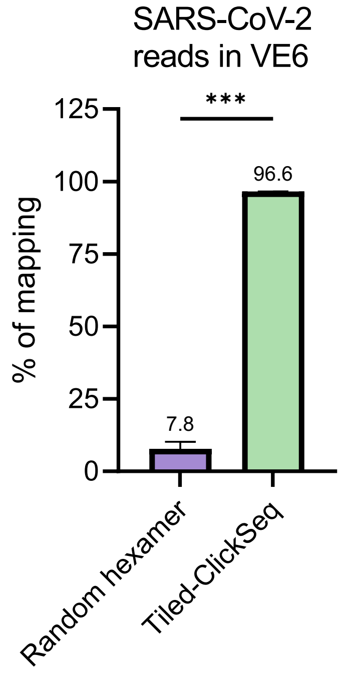


**Extended Data Figure 6.** RNAseq efficiency of Tiled-ClickSeq vs. random hexamer. In complex RNA environment such as cell lysate, Tiled-ClickSeq showed >12-fold higher mapping rate than that of random primer approach (without poly-A enrichment).

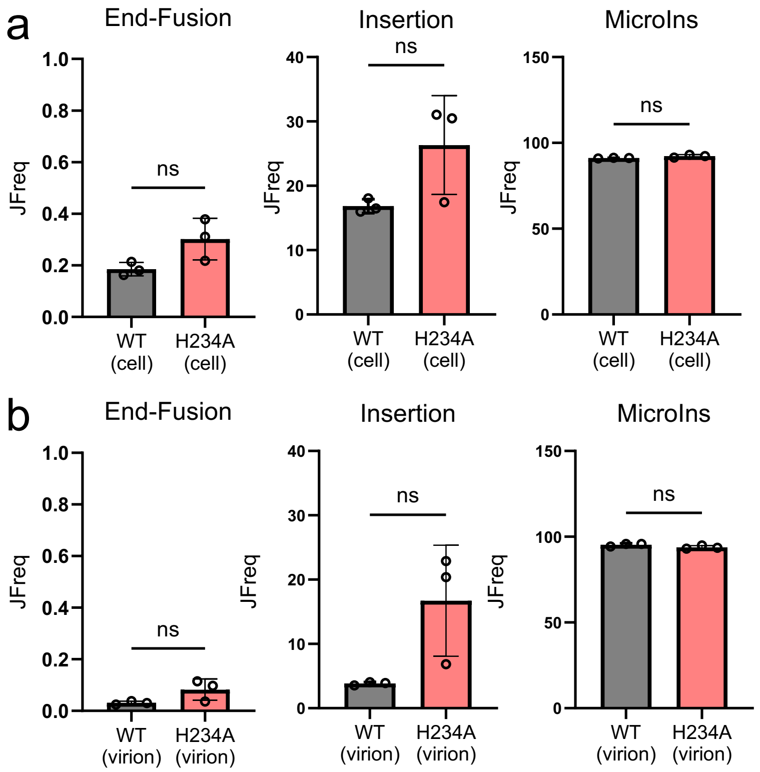


**Extended Data Figure 7.** Other recombination events from SARS-CoV-2 infected Vero E6. In addition to sgmRNA, deletion and MicroDels, we also mapped other recombination events such as end fusions (recombination between 3' end and 5' start of viral RNA genome), insertions (>25nts.) and micro-insertions (microIns, 3-25nts.). No statistical difference was found between WT and H234A in cell lysate (a) or supernatant virions(b).

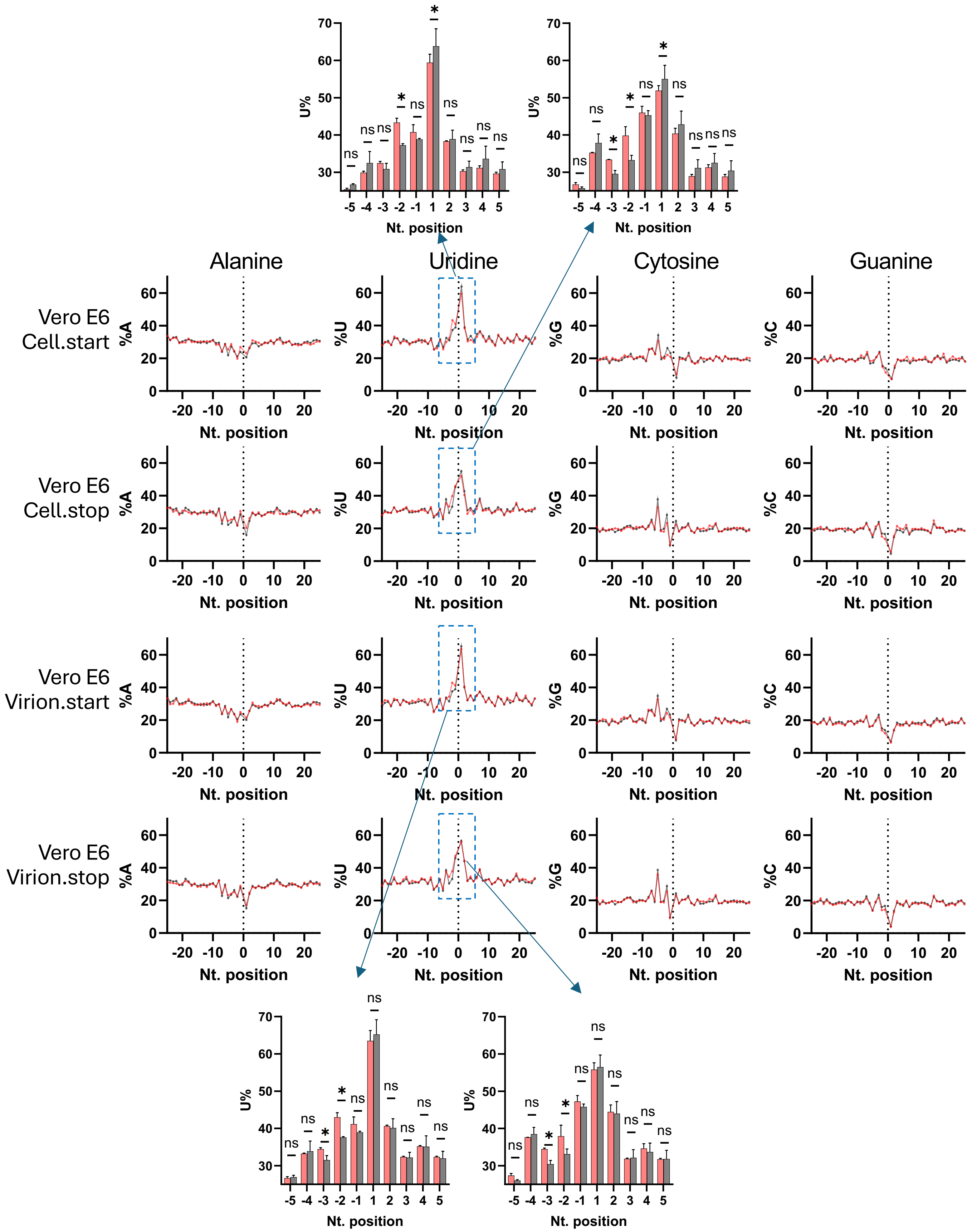


**Extended Data Figure 8.** The frequency of A, U, G, C at each nucleotide position flanking the recombination start (donor) and stop (acceptor) sites in Vero E6 cells infected by WT (black) or H234A (red).

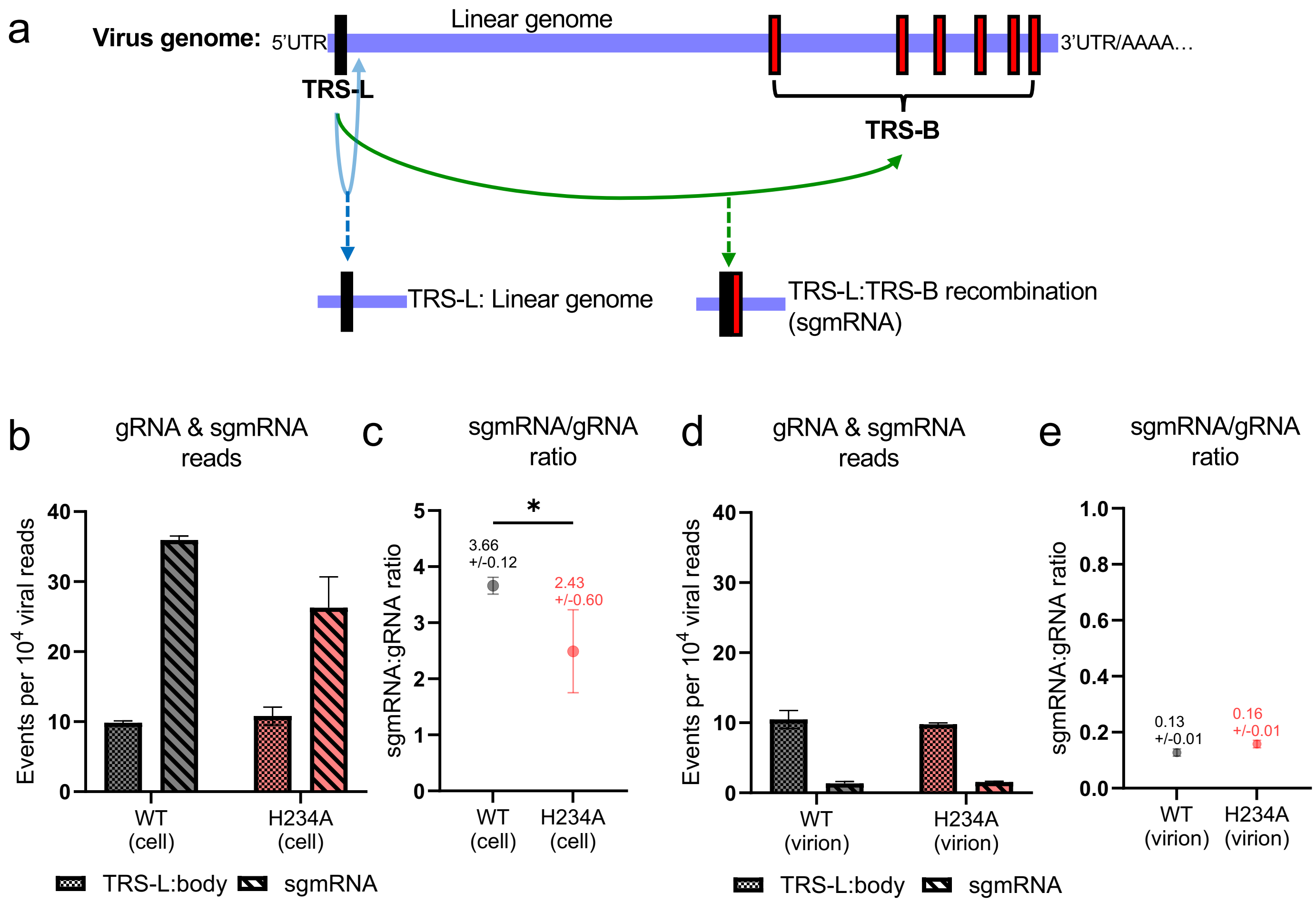


**Extended Data Figure 9.** Genomic RNA to sgmRNA ratio. (a) Schematic of TRS-L:body "junction" ( the "junction" between TRS-L and immediate downstream (nt. 85) of linear genome) and TRS-L:TRS-B junction (recombination between TRS-L of genomic RNA and TRS-B of subgenomic RNA). The JFreq of TRS-L:body remained comparable between WT and H234A in both cell lysate (b) and supernatant (d), but a clear reduction of JFreq of sgmRNA was observed in H234A infected cells (b). The ratio of sgmRNA/TRS-L:body suggests the reduced sgmRNA production in H234A infected cells (c) but not supernatant (e).

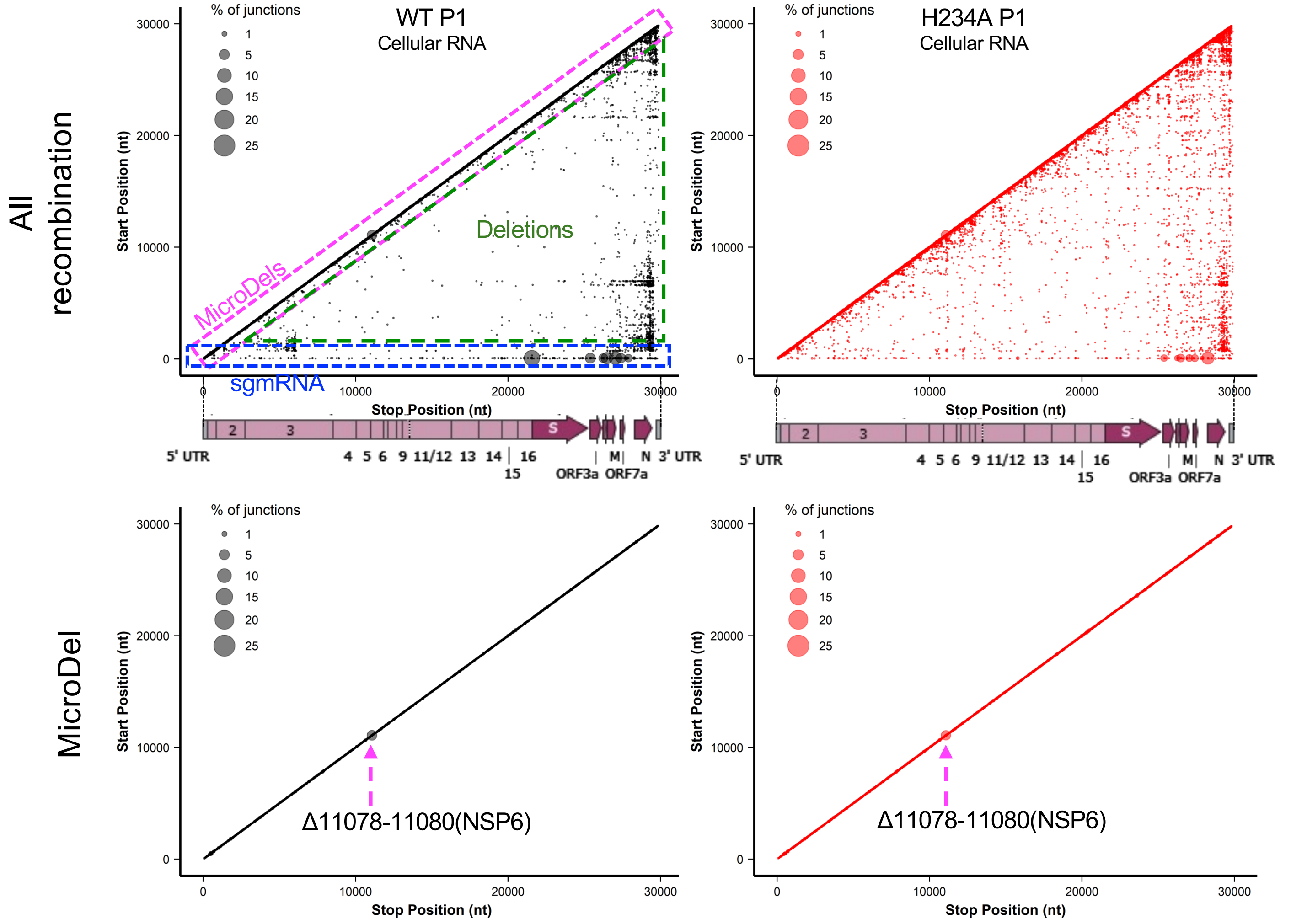


**Extended Data Figure 10.** 2D map of parental strain (P1) all recombination events and micro-deletions. Black = WT, red = NSP15mut. Both WT P1 and H234A P1 contained Δ11078-11080(NSP6) micro-deletion but not the Δ23583-23599(QTQTN) micro-deletion.

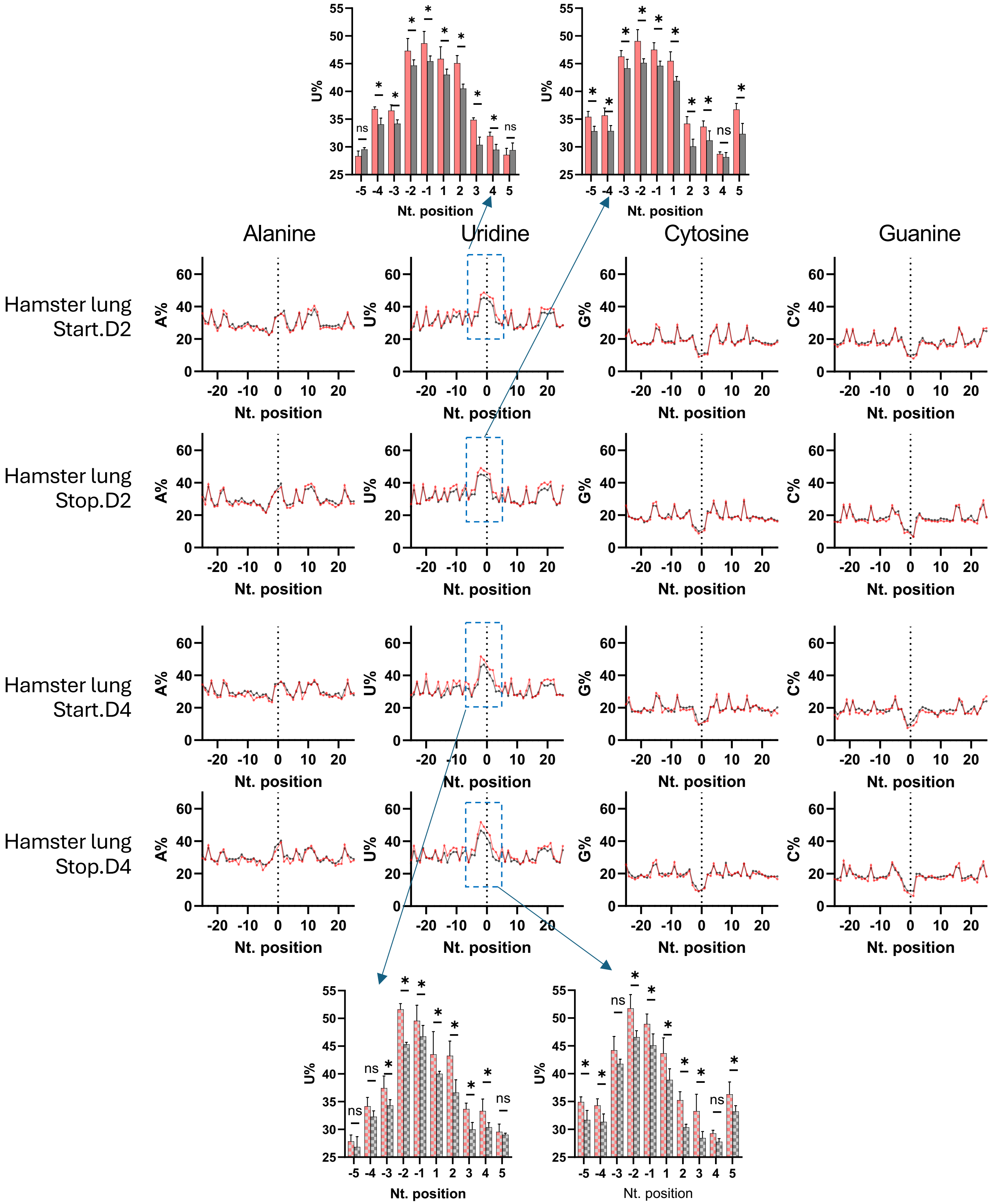


**Extended Data Figure 11.** The frequency of A, U, G, C at each nucleotide position flanking the recombination start (donor) and stop (acceptor) sites in hamster lungs infected by WT (black) or H234A (red).

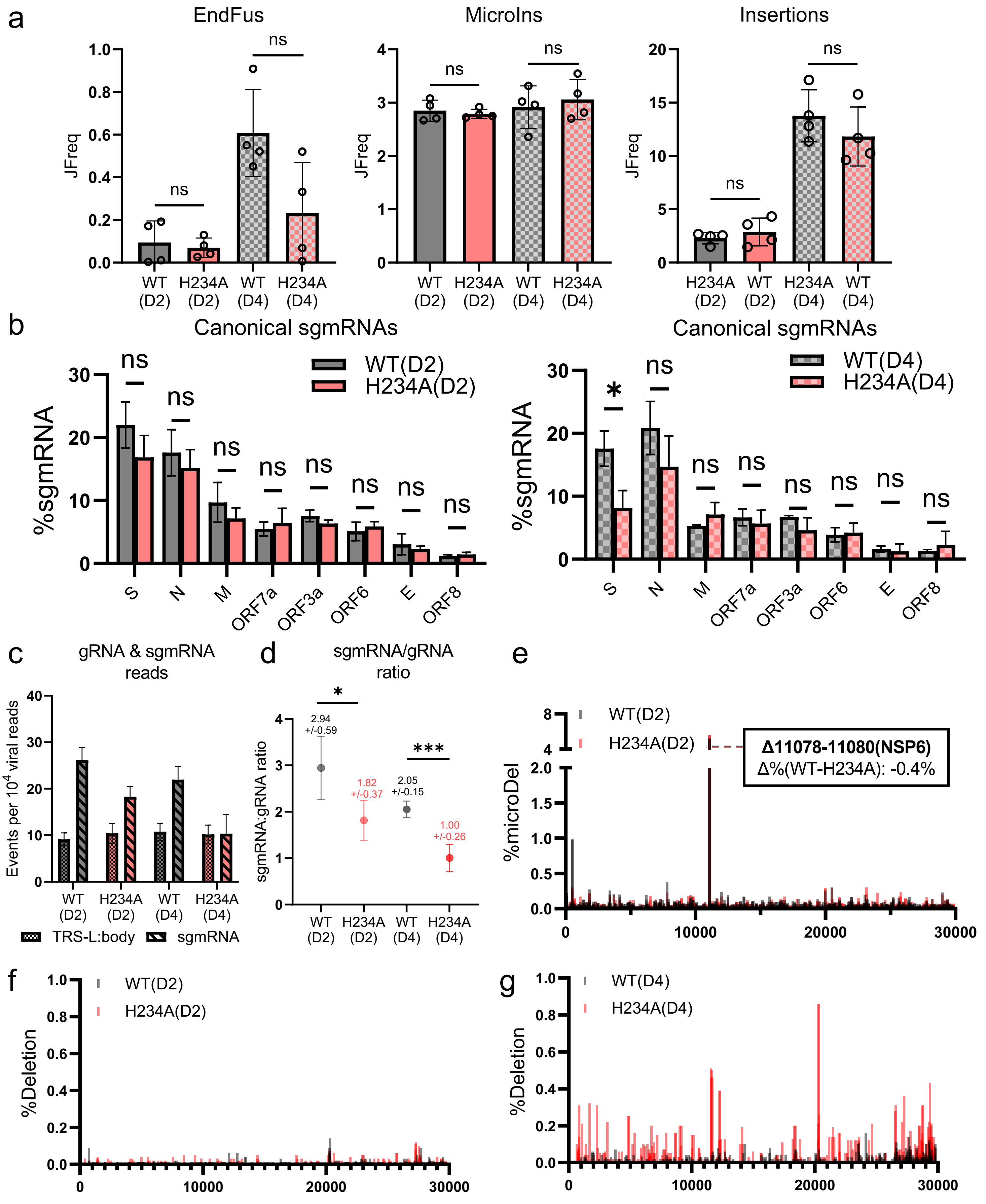


**Extended Data Figure 12. Further characterization of recombination events in hamster lungs.** (a) The JFreq of End-fusions, micro-insertions (3-25nts.) and insertions (>25nts.). No statistical difference was found between H234A and WT at either day 2 (D2) or day 4 (D4). (b) The percentage of canonical sgmRNAs at day 2 and day 4. (c) The frequency of TRS-L:body and sgmRNA. H234A showed clear reduction than WT at both D2 and D4. (d) The ratio of sgmRNA/TRS-L:body. H234A showed significantly reduced sgmRNA production per viral genomic RNA. (e) Micro-deletions of H234A and WT at D2. (f) Deletions of H234A and WT at D2. (g) Deletions of H234A and WT at D4.

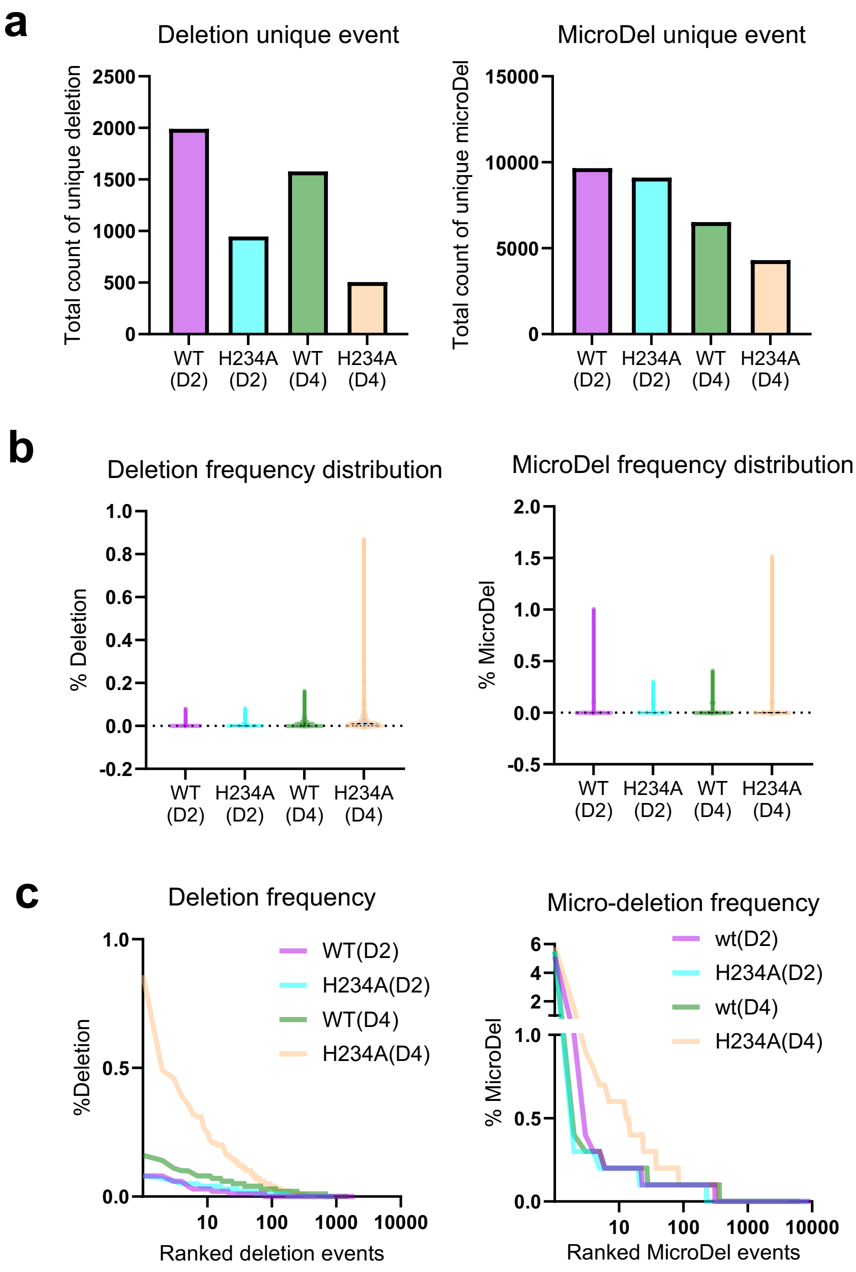


**Extended Data Figure 13.** (a) The total number of unique deletion or micro-deletion events in all replicates. WT-infected animal lungs contained a greater distribution of deletion/micro-deletion events than H234A. (b) The frequency of deletion and micro-deletion events demonstrates that H234A-infected animals accumulated more high frequency deletions/micro-deletions. (c) Elbow plots of deletion/micro-deletion frequency demonstrate that H234A-infected animals contained directed evolution of specific DVG populations.
